# Barley BODYGUARD controls cuticular specialisations regulated by SHINE transcription factors

**DOI:** 10.1101/2025.07.28.666421

**Authors:** Trisha McAllister, Chiara Campoli, Linsan Liu, Tansy Chia, S. Ronan Fisher, Richard Horsnell, Alan R. Prescott, Jennifer Shoesmith, Mhmoud Eskan, Alasdair Iredale, Mirjam Nuter, Luke Ramsay, Micha M. Bayer, Linda Milne, Miriam Schreiber, Vanessa Wahl, Robbie Waugh, James Cockram, Sarah M. McKim

## Abstract

The outer epidermis of land plants secretes a cuticular layer, a hydrophobic diffusion barrier which minimises water loss into the atmosphere and protects from pests, ultraviolet light and organ fusion. Cuticles typically comprise a polyester cutin matrix embedded and overlaid with cuticular waxes, but their exact chemical make-up, structure and functions can vary widely depending on the tissue and species. Barley shows two such cuticular specialisations: (1) deposition of a thick β-diketone-rich wax bloom on multiple organs at reproductive stage, common in other Poeceae species and linked to yield; and, (2) secretion of a sticky layer on the grain fruit (caryopsis) pericarp cuticle which adheres to inner floral hulls, leading to barley’s distinctive ‘covered’ grain used in animal feed and malting. Two SHINE/WAX-INDUCER transcription factors in barley, HvWIN1 and NUD, promote the wax bloom and hull to caryopsis adhesion, respectively, yet little is understood about other genes involved. Leveraging near-isogenic lines of wax-deficient mutants, we identify the barley *BODYGUARD1* (*HvBDG1*) gene encoding an α/β-hydrolase essential for leaf cuticular integrity and wax bloom deposition. Modelling of functional and defective alleles suggests that HvBDG1 N-terminal region control of protein flexibility is important for HvBDG1 function. In addition to their role in controlling barley epicuticular wax deposition, we show that both HvBDG1 and HvWIN1 are essential for strong hull to caryopsis adhesion. Along with NUD, these gene products differentially contribute to ultrastructural changes on the pericarp associated with a cuticular building programme driven by NUD and HvWIN1 regulation of cuticle metabolism and transport and cell wall-related genes, and correlate with shifts in pericarp surface chemistry. We also show that the previously asserted ‘grain-specific’ role of *NUD* should be revised, as our findings reveal that it is essential for maintaining leaf cuticle integrity. Our analyses in barley suggest that NUD and HvWIN1 control cuticular specialisations and cuticle integrity in part via promotion of *HvBDG1* expression, while HvWIN1 and NUD likely act independently from each other. Lastly, mining tetraploid wheat mutant populations followed by crossing to combine mutated homoeologues demonstrated that BDG1 and WIN1 orthologues also control wax bloom in wheat. Taken together, our work greatly expands the genetic networks and molecular activities important for cuticle development in cereals and the underlying mechanisms for both shared and species-specific cuticular specialisations.

## INTRODUCTION

The plant cuticle is a vital hydrophobic diffusion barrier covering almost all aerial surfaces in land plants and protecting from desiccation, pathogen attack and tissue fusion (Ingram and Nawrath, 2017; Arya et al., 2021; González-Valenzuela et al., 2023). Consisting of a cutin matrix embedded with polysaccharides and cuticular waxes, the cuticle is overlaid with epicuticular waxes typically made from very long chain fatty acids (VLCFAs) and their derivatives (Yeats and Rose, 2013). However, within this general framework cuticles show remarkable differences in composition, structure and function. For instance, in contrast the alcohol-rich wax found in wild Triticeae species (Tulloch, 1980), leaf tissues, stems and spikes of the related cereal crops, barley (*Hordeum vulgare* L.) and wheat (*Triticum* spp.), develop glaucous (white-blue) epicuticular wax blooms dominated by β-diketones rather than VLFCAs (Mikkelsen, 1979; Adamski et al., 2013). The expression of these wax blooms is drought responsive and confer improved water use efficiency and ultraviolet light reflectance, suggesting a possible reason for their selection during domestication (Bi et al., 2017; Richards et al., 1986).

Studying cuticle mutants, in particular the *eceriferum* or *cer* (“not bearing wax”) fusion or glossy mutants isolated in multiple species, helped identify several genes encoding metabolic components, transporters, signalling proteins and transcription factors involved in cuticle formation and maintenance, many of which appear broadly conserved (Samuels et al., 2008; Kong et al., 2020). For instance, the WAX-INDUCER/SHINE (SHINE)-like transcription factors (WIN/SHN) are critical for cuticle development in Arabidopsis, rice, wheat and tomato (Shi et al., 2011, 2013; Wang et al., 2012; Zhou et al., 2014; Kannangara et al., 2007; Bi et al., 2018), while the GDS(L) [Gly, Asp, Ser, (Leu)] motif esterase/lipases (Brick et al. 1995) in barley and rice promote cutin accumulation and water retention (Park et al., 2010; Li et al., 2017). In contrast, other genetic components appear more specialised, such as the *CER-CQU* metabolic gene cluster encoding a novel polyketide synthase that generates the β-diketones and hydroxy-β-diketones dominating the glaucous wax bloom in barley and wheat (Hen-Avivi et al., 2016; Schneider et al., 2016). Recently, we revealed that defective alleles in *HvGDSL1* or a *SHN/WIN* homologue, *HvWIN1*, cause reduced wax blooms due to lower *CER-CQU* expression (McAllister et al., 2022; Campoli et al., 2024), demonstrating that conserved cuticle genes can regulate diverse cuticular components.

While barley and wheat both develop wax blooms, barley develops a unique cuticular specialisation on its caryopsis (or grain fruit). During grain development the outer pericarp extrudes an adherent ‘cementing’ layer that sticks to the inner face of the floret hulls, leading to barley’s distinctive ‘covered grain’ phenotype (Gaines et al., 1985). During post-domestication cultivation, deletion of *NUDUM* (*NUD*), another *WIN/SHN* homologue in barley, led to a loss of hull adhesion resulting in naked grain barley subsequently cultivated for human food (Gaines et al., 1985; Taketa et al., 2008). Naked grain lacks the pericarp’s cementing layer, yet *NUD* transcripts reportedly accumulate in the underlying seed coat or testa (Gaines et al., 1985; Taketa et al., 2008). We know very little about the steps between *NUD* expression, downstream targets and pericarp surface changes.

Naked barley for human food use is a growing market, yet most barley is grown for animal feed or brewing, end uses where a strongly adherent hull is desirable to protect grain and increase malting efficiency (Okoro et al., 2017). Grain skinning or partial hull shed of covered barley leads to low quality malt and is an increasing problem for barley growers and end users (Okoro et al., 2017). We observed that a subset of barley *cer* mutants with defective wax blooms also show grain skinning (Campoli et al., 2024) and speculated that these phenotypes could reflect disrupted function in targets downstream of NUD. Here, we reveal that one such *cer* mutant results from defective alleles in the *HvBODYGUARD1* (*HvBDG1*) gene, so named due to homology with the BDG alpha-beta hydrolase necessary for cuticular integrity and accumulation of epicuticular wax and cutin in Arabidopsis (Kurdyukov et al., 2006a; Jakobson et al., 2016). Comparative modelling of wild-type and mutant HvBDG1 proteins suggests that the HvBDG1 N-terminal region controls flexibility important for function while transient heterologous expression shows that HvBDG1 localises to the endoplasmic reticulum (ER) and mobile spherical bodies. Defective alleles of *HvWIN1* also led to grain skinning, while deletion of *NUD* compromises leaf cuticle integrity. These findings, combined with spatial expression patterns, genetic analyses and comparative transcriptomics suggest that HvWIN1, HvBDG1 and NUD have unique and overlapping roles in developing grain and leaves, with HvBDG1 potentially acting downstream of either factor. We also demonstrate that loss of HvBDG1 function causes increased cuticle permeability in barley leaf blades and sheaths, and that BDG1-mediated promotion of wax blooms and cuticle integrity and WIN1 promotion of the wax bloom is conserved in the related crop species durum wheat (*T. turgidum* ssp. *durum*). Our work expands the genetic networks and molecular activities important for cuticle development in barley and our understanding of the mechanisms underlying both shared and species-specific cuticular specialisations in plants.

## MATERIALS and METHODS

### Plant material and growth conditions

Germplasm details are provided in Table S1. Barley cultivar (*cv.*) Bowman and the Bowman Near Isogenic Lines (BW-NILs) for introgressed *cer* alleles (BW156, BW406, BW407 and BW126; Druka et al., 2011) were obtained from the James Hutton Institute (JHI). Original barley mutant alleles and parent *cv*s were obtained from NordGen (Nordic Genetic Resource Center), the U.S. National Plant Germplasm Service (NPGS), and Okayama University, Japan. Barley plants for genotyping and phenotyping were grown in glasshouses under long day photoperiod conditions (16 hr light 18°C/8 hr dark 14°C) in standard cereal compost mix: 1.2m^3^ Peat, 100l sand, 1.5kg Osmocote Exact Start, 3.5kg Osmocote Exact 3-4 month, 2.5kg Lime, Ca and Mg (each), 0.5kg Celcote, 100l Perlite and 280g Intercept. Double mutants were generated by crossing and confirmed by genotyping segregating F_2_ populations. Barley plants for grain skinning and debranner assays were grown in pots in an outdoor polytunnel, supplied with automated watering and disease control when required, at the James Hutton Institute (56.456627, -3.0675811). Plants were watered thrice daily for three minutes each time. Disease control including Opus Team spray for the Mildew in sprayed Opus Team and Hallmark for aphids. The cereal potting mix was as follows: Peat/Wood Fibre 70%/30%, Lime, Nutricote 70 Day 16-10-10, Base 15-10-20+TE, Nitrochalk, Mircomax, Aquasorb 3005, Wetting Agent and Perlite.

### Grain skinning and debranner assays

Mature grain harvested from plants grown during spring and summer of 2014 and 2018 was used to assess grain skinning using manual skinning assays and debranner assays (Campoli et al., 2024), respectively. To measure the proportion of barley grains which skin in a sample, mature grain from each genotype harvested from a single plant for each biological replicate (Bowman, n=1 pool of three plants; BW156, n=4, BW406, n=4 and BW126, n=3) was manually threshed and visually assessed for signs of skinning. Skinning was also assessed from mature grain harvested from a single plant, whereby a 25 g sample of mechanically threshed grain was processed using a Lab Scale Debranner TM05 (Satake Seed Mill) for 15 s, as described in Campoli et al. (2024). Skinning severity was represented as the weight of hull lost compared to total hull weight of the sample. Double mutants were similarly assessed by manual threshing along with Bowman and single mutant parent controls. Results were tested for significance using Analysis of Variance (ANOVA) and Tukey’s HSD post-hoc test.

### Cuticle integrity

Four biological replicates were harvested for each barley genotype by sampling 10 cm sections of fresh tissue from the second fully expanded leaf blade (growth stage 12, GS12; Zadoks et al, 1974) and flag leaf sheaths at GS55. Fresh weight was recorded for each sample and chlorophyll leaching assays subsequently performed as described by McAllister et al. (2022) with one modification: samples were kept in 80% ethanol for up to 72 hours to measure the total chlorophyll leached and hourly measurements were plotted as a percentage of this total. Statistical significance was evaluated by ANOVA and Tukey’s HSD using Area Under the Curve (AUC). Data were plotted in RStudio (Posit team, 2025) using ggplot2 (Wickham, 2016). For wheat leaves, we collected the second leaf blades from 2-week-old plants, trimmed the ends to retain the middle segments, and exposed them to 0.05% Toluidine Blue for five hours at room temperature to assess how quickly the dye penetrates the epidermal surface to stain the underlying cells.

### Electron Microscopy

Barley caryopses were harvested from the middle of the inflorescence at seven and 11 days post anthesis (DPA) (two caryopses harvested from individual plants with five individuals per genotype). Hulls were manually removed from all samples. Caryopsis sections were taken from the centre of the dorsal surface for Scanning Electron Microscopy (SEM) and Transmission Electron Microscopy (TEM). Samples were fixed in 4% formaldehyde and 2% glutaraldehyde in 70 mM PIPES at room temperature for four hours then overnight at 5°C. Samples were washed in 0.1 M sodium cacodylate three times for ten min and stored in 0.1 M sodium cacodylate at 5°C. SEM samples were post-fixed in 1% osmium tetroxide in 0.1 M sodium cacodylate for one hour at room temperature, washed twice for ten min with 0.1 M sodium cacodylate, ethanol dehydrated, critical point dried and sputter coated with 10 nm palladium. Samples were imaged using a JEOL JSM-7400F SEM.

TEM samples were processed as above with the following modifications: 1.5% potassium ferrocyanide was added to the post-fixation step. After washing in 0.1 M sodium cacodylate, samples were washed twice with 0.1 M sodium acetate buffer for ten minutes and stained with 1% uranyl acetate in acetate buffer for one hour. Samples were washed twice in distilled water for ten min, ethanol dehydrated, washed twice for ten min in 100% propylene oxide and incubated in 50% Durcupan resin in propylene oxide overnight at room temperature. Samples were transferred to 100% Durcupan resin for eight hours and embedded in Durcupan resin overnight at 60°C. Sections of 70-100 nm were cut on a Leica Ultracut UCT, stained with 3% uranyl acetate and Reynolds lead citrate for 20 min each and imaged on a JEOL 1200EX TEM using a SIS III camera.

### Lipid chemistry

Surface waxes were extracted by dipping barley caryopses and hulls at five and 11 DPA in 10 mL of dichloromethane (DCM) containing 10 µg of methyl-nonadecanoate as the internal standard. Sample size was nine to 16 caryopses or derived hulls per replicate. Extracts were subsequently dried and derivatised in 80 µL N-O-bis-trimethylsilyltrifluoroacetamide (BSTFA). Wax components were identified using Gas Chromatography–Mass Spectrometry (GC–MS) using a Trace DSQTM II Series Quadrupole system (Thermo Electron Corporation), fitted with a CTC CombiPAL autosampler (CTC Analytics). Run conditions and analysis were as described in Campoli et al. (2024). After surface wax extractions, samples were returned to DCM for exhaustive wax extraction over a period of two to three weeks with frequent change of solvent and dried fully before proceeding with cutin extraction. Cutin extraction, run and analysis was performed as described in Campoli et al. (2024).

### Gene mapping, candidate gene sequencing and whole genome survey sequencing

DNA extracted from Bowman, *cv.* Bonus and BW156 leaf tissue was used for genotyping with the 50K iSelect single nucleotide polymorphism (SNP) array (Bayer et al., 2017) as described in Arrieta et al. (2021) to identify donor introgressions in BW156. High-confidence gene models within the introgression interval of BW156 were identified based on *cv*. Morex V1 (GCA_901482405.1) and V2 genome assembly (GCA_902498975.1) sequence data and gene predictions in BARLEX (Colmsee et al., 2015; Mascher et al., 2017; Monat et al., 2019). Coding sequences (CDS) were confirmed by comparing against gene models predicted from the genome assembly of *cv.* Golden Promise (GCA_902500625.1; Schreiber et al., 2020) as well as comparing with predicted protein sequences from homologous gene models in other species collected using NCBI (Genes, 2004) and TAIR (Berardini et al., 2015). The entire CDS of every high-confidence gene in the introgression was polymerase chain reaction (PCR) amplified from BW156, Bowman and Bonus DNA (primers in Table S2) using either (a) GoTaq® in a final reaction volume of 15 µL: 3 µL 5x Colourless GoTaq® Reaction Buffer (Promega), 0.3 µL PCR Nucleotide mix, 0.3 µL of each 10 µM primer, 1 µL template DNA, 0.075 µL GoTaq® G2 DNA Polymerase (Promega), or (b) Q5® polymerase in a final reaction volume of 25 µL: 5 µL 5x Q5® Reaction Buffer (New England Biolabs, NEB), 0.5 µL PCR Nucleotide mix, 1.25 µL of each 10 µM primer, 1 µL template DNA, 0.25 µL Q5® High-Fidelity DNA Polymerase (NEB). PCR products were cleaned using ExoSAP-IT™ PCR Product Cleanup Reagent (Thermo Fisher Scientific). 2 µL of reagent was added per 5 µL PCR sample and incubated at 37°C for 30 min, 80°C for 30 min. PCR amplicons were sent for Sanger sequencing at Genome Technology Lab at the James Hutton Institute (JHI) and resulting sequencing data was analysed using Geneious 10.2.6 (Biomatters). We PCR amplified and sequenced the *HvBDG1* gene in the original mutant donor for BW156, *cer-z.52,* and eight other *cer-z* alleles. We also sequenced the *HvBDG1* gene in BW156 lines obtained from four different stock sources (seed store at the James Hutton Institute, USDA supplied seed, NordGen supplied seed, and a lab stock bulked in 2019), and in the *cer-a* BW-NIL BW406 and 65 *cer-a* alleles, along with their respective parental lines (Table S1).

Whole-genome sequencing of genomic DNA extracted from BW156, Bowman and Bonus was performed on the Illumina NovaSeq platform. Raw reads were mapped and variants called as described in Liu et al. (2022). We retained only variants containing predicted missense mutations in high-confidence models predicted in Morex V2 genes in the mutant compared to Bonus and Bowman.

### Gene expression analyses

We mined existing barley expression data for *HvBDG1* on the EORNA database (Milne et al., 2021) searching by transcript model and exporting normalised expression. For expression analysis by quantitative reverse transcription-PCR (qRT-PCR), we harvested Bowman caryopses at 3, 5, 7, 9 and 11 DPA and separated these from the palea and lemma. A minimum of three caryopses were harvested per individual, pooling three individuals in each of three biological replicates per timepoint. Samples were snap-frozen in liquid nitrogen and stored at −70°C. RNA was isolated using TRI Reagent (Sigma) as described in Patil et al. (2019). cDNA was synthesised from each sample using the ProtoScript II First Strand cDNA Synthesis Kit in a reaction containing 1 µg RNA (NEB). qRT-PCR was performed using Roche FastStart Universal Probe Master (ROX) and gene-specific primers (Table S2) for *HvBDG1*, with *PPA* (*HORVU.MOREX.r3.5HG0522310*) and 26S (*HORVU.MOREX.r3.7HG0714100*) genes used as endogenous controls. Each cDNA sample was diluted to 1:3 with dH_2_O for the template. Each reaction consisted of 5 µL template and 20 µL reaction mix (12.5 µL FastStart, 0.25 µL probe, 0.25 µL each primer, 6.75 µL dH_2_O. We also harvested four biological replicates of caryopses at 5 and 9 DPA (pooling three and four individuals per biological replicate, respectively) from Bowman, BW638 and BW407, leaf sheath tissue (pooling six individuals per biological replicate) from Bowman, BW156 and BW407 at GS39 as in Liu et al. (2022), and leaf blade tissue (pooling five individuals per biological replicate) from Bowman, BW638, BW407 and BW638/BW407 double mutant at GS11. Two sections of leaf blade were harvested: a mid-section containing 3 cm below the point of leaf emergence and a basal section containing the remaining portion at the base of the leaf. RNA was isolated using the QIAGEN RNeasy Kit as in McAllister et al. (2022) and cDNA was synthesised using SuperScript IV VILO Master Mix (Invitrogen) in a reaction containing 2.5 µg RNA for grain and leaf blade, and 2 µg RNA for leaf sheath. qRT-PCR was performed using SYBR Green PowerUp (Thermo Fisher Scientific) and gene specific primers for *HvBDG1*, with 3*-PHOSPHOINOSITIDE-DEPENDENT PROTEIN KINASE 1* (*PDPK1*, *HORVU7Hr1G096480*) and *Actin* (*HORVU5Hr1G039850*) genes used as endogenous controls for leaf blade samples, and *PDPK1* and *TRICHOME BIREFRINGENCE-LIKE 25* (*TBL25*, *HORVU1Hr1G029950; HORVU.MOREX.r3.1HG0031980*) genes as endogenous controls for grain samples, as well as *HvCER-U*, with *Actin* as the endogenous control in leaf sheath samples. Templates were prepared by diluting each cDNA sample with dH_2_O to 1:3 for leaf blade samples, 1:2 for grain samples, and 1:2.5 for sheath samples. Reactions consisted of 4 µL template and 16 µL reaction mix: 10 µL SYBR Green PowerUp, 0.3 µL each primer (20 µM), 5.4 µL dH_2_O. All qRT-PCR reactions were run on a StepOnePlus (Applied Biosystems) according to manufacturer’s instructions with three technical replicates per sample, three biological replicates per genotype and a no-template control per target on each plate. Relative expression was calculated using the 2^−ΔΔCt^ method (Pfaffl, 2001).

For gene expression analyses using *in situ* hybridization, caryopses at 5 and 9 DPA were harvested and fixed in 4% formaldehyde in PEM Buffer [0.1 M PIPES (pH 6.95), 1 mM EGTA, 1 mM MgSO_4_]. Tissues were processed in a Leica TP1020 automated tissue processor with the following steps: 1 h in 70% ethanol, 1 h 30 min in 80% ethanol, 2 h in 90% ethanol, 1 h in 100% ethanol, 1 h 30 min in 100% ethanol, 2 h in 100% ethanol, 1 h in 100% xylene, 1 h 30 min in 100% xylene, 2h in Paraplast® wax at 65°C and a final step of 4 h in Paraplast® wax at 65°C. Samples were embedded using a Leica EG1160 wax embedder and sectioned using a Leica RM2265 microtome at 8 μm thickness. Probe synthesis, slide processing and *in situ* hybridization were done as described by Gramma and Wahl (2023). Regions of *HvBDG1* (370 bp) and *HvWIN1* (400 bp) transcripts were PCR amplified from Bowman cDNA using Q5® polymerase and cloned into pCR-Blunt (Thermo Fisher Scientific) and pMini-T 2.0 (NEB), respectively, in antisense orientation from the T7 promoter. Antisense templates including the T7 promoter were amplified from plasmid DNA for *HvBDG1* and *HvWIN1* (BDG AS plasmid, BDG_C2_AS; WIN1 AS plasmid, WIN1_C2_AS) using GoTaq G2® polymerase and M13 primers for *HvBDG1* and using WIN1_IS_F2, WIN1_IS_R2 primers for *HvWIN1*. A *NUD* fragment (291 bp) was PCR amplified from Bowman cDNA using GoTaq G2® polymerase (Promega) and primers with an added T7 binding site to generate the antisense template. Sense templates were made as described in Campoli et al. (2024). All templates were phenol chloroform purified prior to probe synthesis. Digoxigenin-labelled sense and antisense probes were synthesised using T7 polymerase according to the DIG RNA labelling kit (SP6/T7) (Roche) instructions from purified templates. Primer sequences for probe template generation are listed in Table S2. Slides were imaged using a Zeiss Axioscan 7 Slide Scanner at 20× magnification using brightfield transmitted light by the Dundee Imaging Facility.

### Phylogenetic analyses

To identify BDG homologues in model plant species, HvBDG1 was used as a query for BLASTp searches against predicted protein databases held at Ensembl Plants (https://plants.ensembl.org/), FernBase (https://www.fernbase.org/), SolGenomics (https://solgenomics.net/), Phytozome (https://phytozome-next.jgi.doe.gov/) and PLAZA Gymnosperms (https://bioinformatics.psb.ugent.be/plaza/versions/gymno-plaza/). Protein sequences were checked by comparison of the corresponding *BDG* genomic sequence with that of other *BDG* homologues and manually corrected where necessary. Full predicted protein sequences were aligned using Geneious 10.2.6. and Molecular Evolutionary Genetics Analysis 11 (MEGA11) (Tamura et al., 2021) using MUSCLE (Edgar, 2004). Evolutionary history was inferred using the Maximum Likelihood method and JTT matrix-based model in MEGA11 (Jones et al., 1992) and tested using 500 bootstrap replications. Full protein sequence alignments were examined for conserved motifs amongst angiosperm BDG proteins. Sequences were designated as motifs if at least two residues were 100% conserved within the alignment.

We analysed relationships with BDG proteins across the green plant lineage to examine BDG protein emergence in land plant evolution. BDG sequences from *Chlamydomonas reinhardtii*, *Chara braunii*, *Marchantia polymorpha*, *Physcomitrium patens* and *Amborella trichopoda* were profiled from Ensembl Plants (https://plants.ensembl.org/) and sequences from *Klebsormidium nitens* profiled from the *Klebsormidium nitens* NIES-2285 genome project (www.plantmorphogenesis.bio.titech.ac.jp/).

### Protein structure analyses

HvBDG1 protein secondary structure was initially predicted using Phyre 2 in intensive mode (Kelley et al., 2015). Protein sequences for HvBDG1 (GenBank acc: KAI4977554.1) and the H407R mutant were submitted to the NCBI Conserved Domain Database (Marchler-Bauer et al., 2015; 2017), InterProScan v98.0 (Paysan-Lafosse et al., 2023) and CDVist (Adebali et al., 2015) for the identification of conserved domains. Structural predictions were conducted using the Integrated Fold Recognition – Tertiary Structure (IntFOLD-TS) tool hosted by the IntFOLD7 server (McGuffin et al., 2019, 2023) which integrates template-based modelling, *ab initio* and *de novo* modelling dependant on the availability of templates utilising LocalColabFold 1.0.0, a local version of AlphaFold2 (Jumper et al., 2021; Mirdita et al., 2022) and trRosetta2 (Anishchenko et al., 2021) to predict secondary and tertiary structures. Structures were further refined using ReFOLD3 (Adiyaman and McGuffin, 2021) which conducts model quality assessment for error localisation. Refined structures were then assessed using ModFOLD9 (McGuffin et al., 2021), followed by independent quality assessment with DeepUMQA-X (Guo et al, 2022) the stereochemistry of the prediction by ProCheck (Laskowski et al., 2018), and structural flexibility determined using the CABS-flex 2.0 server (Kuriata et al., 2018) with the following settings: Mode: SS2, Gap: 3, Max: 8.0 Min: 3.8. Conservation scores were calculated with ConSurf (Ashkenazy et al., 2016), based on multiple sequence alignments of homologues identified via HMMER against UniRef90 (E-value 1e^-3^; identity threshold: 35-95%), aligned using MAFFT. Structural homologues were identified using the FoldSeek (Van Kempen et al., 2023) and validated through TM-alignment via the RCSB-PDB pairwise structure alignment tool (Bittrich et al., 2024). All structural visualisations were rendered in PyMOL (Schrödinger LLC).

### Cellular localisation

To show HvBDG1 localisation, we used heterologous expression in *Nicotiana benthamiana.* The CDS of *HvBDG1* in Bowman was cloned into pK7WGR2 to generate a p35S driven construct N-terminally fused with Red Fluorescent Protein (RFP). *Agrobacterium tumefaciens* strain GV3101 was transformed with the recombinant binary vector and cultures carrying the BDG-RFP construct and p19 silencing suppressor at an OD_600_ of 0.01 in infiltration buffer (10 mM MES, 10 mM MgCl_2_ and 0.2 mM acetosyringone) were used to infiltrate the youngest fully expanded leaves of approximately four-week-old *N. benthamiana*. *N. benthamiana* plants were grown at 20°C under long day photoperiod (16 hr light/8 hr dark). Leaves were harvested either 48 or 72 hours post infiltration, infiltrated with water to displace air in the samples and mounted on microscope slides using double-sided tape. Abaxial epidermal cells were imaged using a Nikon A1R confocal laser scanning microscope. GFP and RFP were excited at 488nm and 561nm, with emissions at 500-530 nm and 570-620 nm, respectively. To assess co-localisation with cell compartment markers, BDG-RFP was transiently expressed in *N*. *benthamiana* lines expressing CB28 (ER-HDEL-GFP, described in Ruiz et al., 1998) and Lti-PM-GFP (construct described in Kurup et al., 2005 and transformed into a stable line as described in Wang et al., 2017). To further assess co-localisation with other cell compartments, BDG-RFP was co-infiltrated with constructs expressing a Golgi body marker (ST-YFP, construct described in Boevink et al., 1998) and two endosome markers (Ara6-YFP and Ara7-YFP, constructs described in Ueda et al., 2001).

### Transcriptomics

RNA isolated from Bowman, BW638 and BW407 caryopses at 5 and 9 DPA was quality assessed using a Bioanalyzer 2100 (Agilent Technologies). Library construction, sequencing, data pre-processing and differential expression analysis were performed as in Liu et al. (2022) with an expression fold change cut-off of log_2_ 0.5 used to define differentially expressed genes (DEGs).

#### Wheat *BDG1* and *WIN1* mutant identification, germplasm, genetic marker development and phenotyping

To identify wheat orthologues of *HvBDG1* and *HvWIN1*, the barley CDS from gene model transcript *HORVU.MOREX.r3.7HG0644300.1* (*HvBDG1*) and *HORVU.MOREX.r3.6HG0578240* (*HvWIN1*) were used for BLASTn searches of the reference genome of durum wheat (*T. turgidum* ssp. *durum* cv. Svevo, assembly Svevo.v1; Maccaferri et al., 2019) using Ensembl Plants release 60 (Harrison et al., 2024). Exploiting the ethyl methanesulfonate (EMS) mutated Targeting Induced Local Lesions IN Genomes (TILLING) library and associated exome capture sequence data previously generated for tetraploid durum wheat cv. Kronos (2n = 4X = 28, consisting of A and B sub-genomes) (Krasileva et al., 2017), durum mutants with premature stop or splice site mutations predicted to severely affect TdBDG1 or TdWIN1 homoeologue protein function were identified bioinformatically using Ensembl Plants, followed by manual inspection. Seed of TILLING lines were obtained from the SeedStor genebank (https://www.seedstor.ac.uk/) and grown using ‘Niab cereals soil mix’ supplied by ICL (https://icl-growingsolutions.com/) in a naturally lit glasshouse supplemented with Sunblast LED lights (https://kroptek.com/) under a 16h day/20°C, 8h day/15°C regime. DNA was extracted from seedling leaves as described by Fulton et al. (1995). The targeted mutations in the selected *TdBDG1* and *TdWIN1* TILLING lines and derived germplasm were confirmed and tracked using codominant PCR-based Kompetitive allele-specific (KASP) assays. KASP genotyping was done following manufacturer’s instructions (https://3crbio.com/) using the primers listed in Table S2. For each target gene, *TdBDG1* and *TdWIN1*, the TILLING mutations on the A and B sub-genome homoeologues were brought together into the same genetic background via crosses between plants shown by KASP genotyping to be homozygous for the mutant allele. Plants assigned as the female parent were emasculated at the green anther stage, followed by transfer of pollen from plants assigned as the male parent once the stigma of the female parent reached maturity (typically three days after emasculation). The resulting F_1_ grains were grown and self-pollinated to generate F_2_ seed segregating for wild type versus mutant genotype at both homoeologues. Using the KASP assays described above, F_2_ individuals were identified in the following combinations: (i) homozygous for the target mutation in both homoeologues, (ii) homozygous for the target mutation in just one homoeologue and homozygous wild-type for the other homoeologue, or (iii) homozygous wild-type at both homoeologues. Observations of leaf waxiness were made on F_2_ plants, grown in the glasshouse as described above, at the grain filling growth stage.

## RESULTS

### Variation in *HvBDG1* causes defective wax blooms

To learn more about cuticular specialisations in barley, we studied several *cer-z* alleles reported to reduce the wax bloom (Lundqvist and von Wettstein, 1962). We confirmed that BW-NIL156 (BW156), a Bowman near-isogenic line reportedly derived from the neutron radiation-induced *cer-z.52* allele in *cv*. Bonus (Lundqvist and Franckowiak, 1997b), exhibited glossy rather than glaucous leaf sheaths and spikes indicating loss of cuticular wax (Fig 1a). We also observed that BW156 leaf blades and sheaths showed a faster rate of chlorophyll leaching in detached tissue assays compared to Bowman (p = 0.035, Fig 1b and p =1.93 x 10^-5^, Fig S1b, respectively) suggesting cuticular integrity defects. To identify the underlying gene, we genotyped BW156, Bowman and Bonus with the barley 50k iSelect SNP array (Bayer et al., 2017), identifying a single 898 Kbp Bonus introgression on the short arm of chromosome 7 (7HS) in BW156 between markers JHI-Hv50k-2016-451524 (17,533,728 bp) and JHI-Hv50k-2016- 451706 (18,431,796 bp) (Fig S2; Table S3). We initially detected 18 high-confidence (HC) genes within this interval based on the Morex V1 assembly (Mascher et al., 2017). However, comparison to the Golden Promise genome (Schreiber et al., 2020) highlighted 17 gene models with errors including seven false gene models, reducing the list to 11 HC genes. Further cross-checking against the Morex V2 assembly (Monat et al., 2019) identified five new gene models, increasing the total to 16 HC genes in the interval. Sequencing these 16 HC genes identified variation unique to BW156 in only one gene - *HORVU.MOREX.r3.7HG0644300* – which we name *HvBODYGUARD1* (*HvBDG1*) due to homology with *AtBDG1* (AT1G64670), a gene previously shown to control cuticular features in Arabidopsis (Kurdyukov et al., 2006; Jakobson et al., 2016). Compared to Bonus, the *HvBDG1* gene in BW156 has a single non-synonymous A to G polymorphism predicted to change residue 407 in the 501 amino acid HvBDG1 protein from a histidine to an arginine, H407/R (Fig 1c), in the C-terminal region of HvBDG1. Mapping whole-genome sequencing reads from BW156, Bowman and Bonus to the Morex V2 genome assembly, followed by small (< 50 bp) and structural (≥ 50 bp) variant calling, identified the same A to G SNP in the *HvBDG1* gene (Table S4). Another *cer* BW-NIL, BW406, a *cer-a* introgression line derived from the radiation-induced mutant *glossy sheath3.i* (*gsh3.i*) generated in *cv*. Mars (Lundqvist and Franckowiak, 1997a), phenocopied the reduced wax of BW156 (Table S4) and showed an overlapping introgression with BW156 based on 50k iSelect SNP genotyping of BW406 and Mars (Table S2). Sequencing *HvBDG1* in BW406 identified the same A to G SNP (Fig 1c) as in BW156; in fact, of the 64 original *cer-a* alleles available, we identified 60 with mutations in *HvBDG1* (Fig 1c, Table S1): 50 severe disruptions associated with glossy phenotypes, and ten with less disruptive effects associated with intermediate phenotypes (Table S4). In total, we identified 60 independent *cer-a* alleles representing 41 unique variants within *HvBDG1*, suggesting that variation at *HvBDG1* underlies *cer-a*. We detected no change in either the *HvBDG1* CDS or 1400 bp upstream of *HvBDG1* in the nine *cer-z* allelic lines, including the original *cer-z.52*, compared to their respective backgrounds (data not shown). We next sequenced individual plants grown from independent seed stocks from our own in-house seed, seed held by the James Hutton Institute, seed provided from NordGen and USDA-ARS. All showed identical *HvBDG1* variation in BW156 (Table S4). We suggest that BW156 contains an introgressed *cer-a*/*gsh3.i* allele, rather than *cer-z.52*, and that variation in *HvBDG1* underlies *cer-a* alleles. In this manuscript, we subsequently refer to BW156 as *hvbdg1^156^* and BW406 as *hvbdg1^406^* and use Bowman as the wild-type when comparing across these lines.

**Figure 1.**
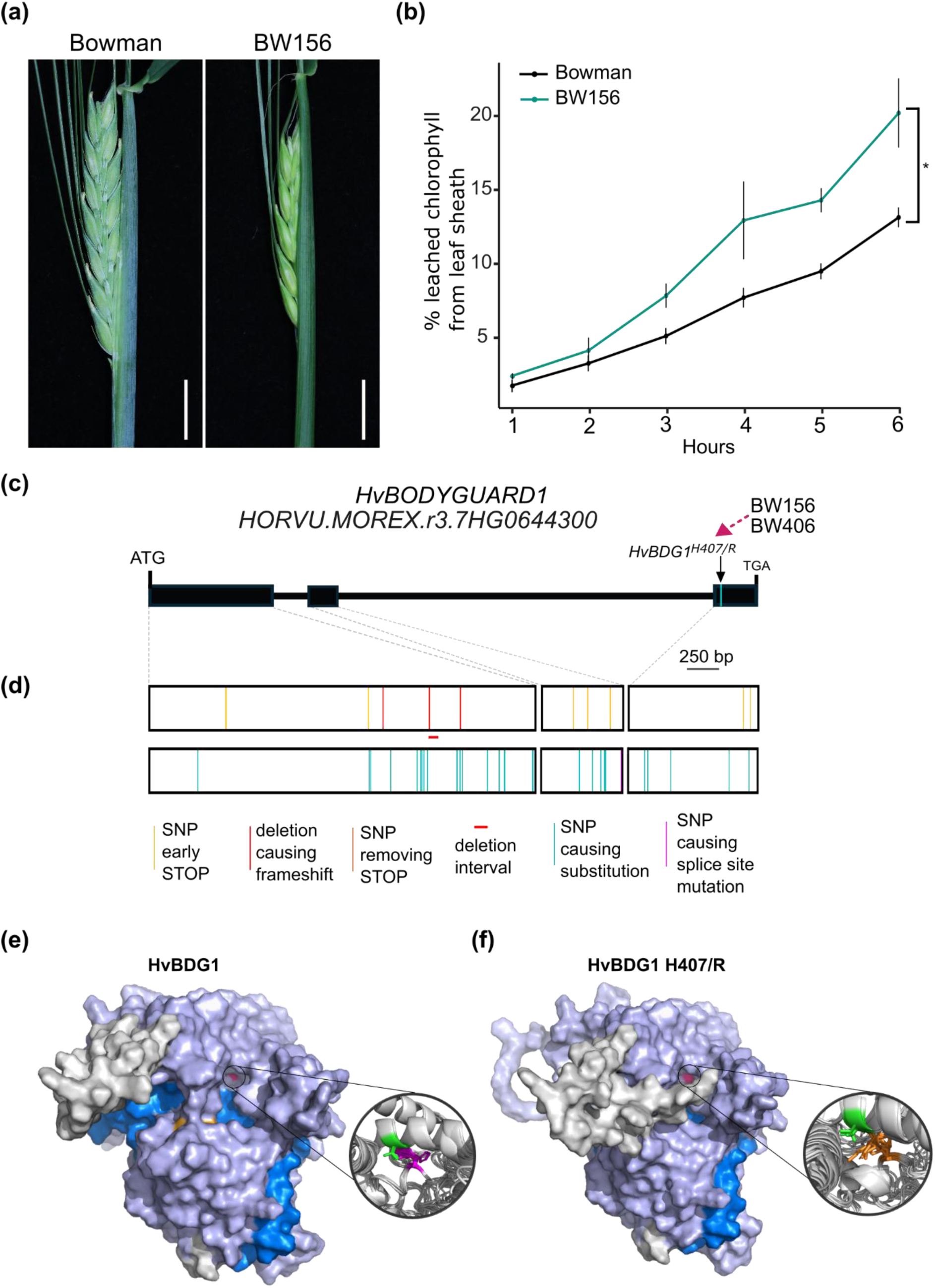
Cuticle defects result from defective *HvBDG1* alleles. (a) Bowman and BW156/*hvbdg1^156^* leaf sheaths. Scale bars, 1 cm. (b) Chlorophyll leaching of detached leaf sheaths from Bowman (n = 3) and BW156/*hvbdg1^156^* (n = 2). (c) *HvBDG1* gene model. Boxes represent exons, horizontal lines represent introns. Gene model represents the full genomic sequence from the start codon to the stop codon. Vertical turquoise line represents A>G SNP found in *hvbdg1^156^* and *hvbdg1^406^* and original mutants which leads to a H4707/R substitution. (d) Boxes represent *HvBDG1* exons with coloured vertical lines representing changes to the HvBDG1 protein predicted from *cer-a* alleles. Top panel: vertical yellow lines indicate SNPs generating early stop codons, vertical red lines indicate deletions causing frameshifts, the horizontal red line indicates a deletion interval, and a vertical orange line refers to a SNP removing the stop codon. Bottom panel: vertical turquoise lines indicate SNPs causing amino acid substitutions and vertical pink lines indicate SNPs causing splice site mutations. Scale bar: 250 bp. (e,f) HvBDG1 protein models where regions and domains are overlaid with an opaque surface. (e) Bonus cultivar (parental wild type) and (f) HvBDG1 H407/R as found in *hvbdg1^156^*. The alpha-beta hydrolase_1 domain shown in dark blue within a larger BDG1 domain containing hydrolase in pale blue. Four active site residues are situated at H225, S299, D448 and H476, shown in orange where S299, D448 and H476 form a catalytic triad and H225 is predicted to stabilise the interaction. Circle insets show H407 interacting with S368 in Bonus HvBDG1 while R407 does not interact with S368 in the H407/R mutant version. As a result, there are no longer any interactions between α-helix 18 and 21.

The glossy appearance and compromised cuticular integrity of *hvwin1^407^*, a BW-NIL in *HvWIN1*, closely resemble *hvbdg1^156^*, although *hvwin1^407^* also has glossy nodes (McAllister et al., 2022; Fig S1b). We detected little expression of the *CER-U* gene in mid-section leaf sheaths in *hvbdg1^156^* and *hvwin1^407^* compared to robust expression in Bowman (Fig S3), consistent with the defective wax blooms in these mutants, indicating that HvWIN1 and HvBDG1 may be essential for normal β-diketone metabolic gene cluster expression. As a transcription factor, HvWIN1 may directly induce *CER-U*, but it is less clear how HvBDG1 regulates metabolic gene expression.

### BDG gene family and protein modelling

We detected two other *BDG*-like genes in barley, *HORVU.MOREX.r3.1HG0051940* and *HORVU.MOREX.r3.2HG0111190*, which we named *HvBDG2* and *HvBDG4*, respectively, based on phylogenetic grouping of their corresponding proteins across selected angiosperm model species (Fig S4). HvBDG1 falls in a grass-specific clade while HvBDG2 sits in a separate grass-specific clade (Fig S4), which may indicate that a duplication giving rise to BDG1-like and BDG2-like proteins occurred after the divergence between monocot and dicot lineages. In contrast, HvBDG4 groups in a clade containing both monocot and dicot members (Fig S4a). Mining the barley gene and transcript expression database, EoRNA (Milne et al., 2021), revealed distinct *HvBDG* expression patterns, although *HvBDG1* was most highly and broadly expressed, with maximal expression in epidermal cells and expanding floral organs, agreeing with its cuticular phenotype (Fig S5). Expanding our phylogenetic analyses across exemplar land plants identified BDG proteins in all species examined, as well as a BDG protein in the terrestrial charophyte *Klebsormidium nitens*, but not in the chlorophyte algae *Chlamydomonas reinhardtii* or the charophyte *Chara braunii* (Fig S4b).

To understand more about BDG protein diversity and how HvBDG1 may work, we aligned selected model angiosperm BDG homologues (Fig S6c). All BDG proteins, including HvBDG1, showed the five highly conserved sequence motifs in the α/β hydrolase domain characteristic of the α/β hydrolase superfamily (Shaw et al., 2002): the oxyanion hole, type II β-turn, a nucleophile elbow containing a catalytic serine and two additional catalytic residues—an aspartate (acidic) and a histidine (basic) (Fig S6; motifs 6, 7, 8, 15 and 16, respectively; Table S5). BDG proteins also contained six completely conserved, novel sequence motifs within the α/β hydrolase domain, of which two corresponded to the structurally conserved α-helix (motif 9) and β-sheet (motif 10) following the nucleophilic elbow and four were situated in a potential lid domain (motifs 11-14). A further four motifs were identified within the N-terminal BDG domain (Fig S6; motifs 2-5), including a transmembrane domain predicted through Phyre2 (Powell et al., 2025) which positions an N-terminal signal peptide in the apoplast. Many *cer-a* amino acid substitution mutations occur within the α/β hydrolase domain or the BDG domain, including the *hvbdg1^156^* allele H407/R substitution (Table S4; Fig S6). Interestingly, HvBDG1 and all its cereal-specific clade members have a unique, highly conserved 11-14 amino acid N-terminal motif (motif 1) absent in other BDGs (Fig S6). Structural models of the *HvBDG1* and *HvBDG1^156^* H407/R variants were generated using the IntFOLD7 server (McGuffin et al., 2019, 2023), which integrates multiple prediction methods as part of IntFOLD-TS including LocalColabFold (a local implementation of AlphaFold2; Jumper et al., 2021; Mirdita et al., 2022), and trRosetta2 (Anishchenko et al., 2021). Model quality was evaluated using ModFOLD9 and independently validated with DeepUMQA-X (Guo et al., 2022) and ProCheck (Laskowski et al., 2018). Model metrics, including Ramachandran plot analyses and steric favourability scores, are provided in Supplementary Note 1 (Fig 1e; Fig S7). The Bonus HvBDG1 model exhibits a canonical α/β hydrolase core composed of a central β-sheet flanked by α-helices and a catalytic triad, comprising S299, D448 and H476, as well as a V-shaped lid domain made up of four alpha-helices arranged over a moderately positively charged active site nestled within a hydrophobic pocket. The HvBDG1^156^ H407/R substitution leads to a histidine-threonine-arginine disrupting a conserved histidine-threonine-histidine which lies in a buried and evolutionarily conserved region with predicted structural importance based on ConSurf analysis (Fig S7). Coarse-grained protein modelling showed that the H407 residue resides buried beyond the Nʹ terminus of α- helix 21 where it is stabilised by an interaction internal to α-helix 18 with S368 (Fig S7). In contrast, the substituted arginine residue in the HvBDG1^156^ H407/R protein is not predicted to interact with S368 and consequently, α-helices 18 and 21 would not stabilise each other, leading to a local conformational rearrangement (Fig 1f) due to increased flexibility in the preceding 150 aa and decrease in flexibility in between 375-400 aa (Fig S7). These structural flexibility changes may lead to a compressed local area and lid conformation in HvBDG1^156^ H407/R, which we speculate could obstruct substrate access to the active site, consistent with the *hvbdg1^156^* allele phenocopying deletion alleles (Table S1).

### Cellular localisation of HvBDG1

Unlike HvWIN1 which likely functions by regulating target gene expression, the link between the predicted alpha-beta hydrolase activity of HvBDG1 and cuticle regulation is less clear. Kurdyukov and colleagues (2006a) reported an extracellular localisation for AtBDG1 and hypothesised a role in extracellular cutin polymerisation. We also explored whether localisation of HvBDG1 could provide hints to its molecular mechanism. Transient expression of an N-terminally tagged red fluorescent protein (RFP)-HvBDG1 protein in *N*. *benthamiana* revealed fluorescence within reticulate structures, a feature characteristic of the ER, and in mobile spherical bodies (Fig 2a,b; <u>VIDEO1</u>). Transient expression of RFP-HvBDG1 in a line expressing an ER-GFP lumen marker (Haseloff et al., 1997; Ruiz et al, 1998), showed overlap in RFP and GFP signals (Fig 2c), although not in the mobile spherical bodies. Transient expression of RFP-HvBDG1 in a line expressing the plasma membrane localised Lti-PM-GFP marker (Kurup et al., 2005) showed possible overlap in small regions which could reflect RFP-HvBDG1 localisation to the plasma membrane or within the ER at direct contact sites between the ER and the plasma membrane (Fig 2d). We also co-expressed RFP-HvBDG1 with the Golgi body marker ST-YFP which tethers YFP to the Golgi membrane by a transmembrane domain (Boevink et al., 2002) as well as with YFP-tagged endosome markers, Ara6 and Ara7 (Ueda et al., 2001). None of these markers co-localised with RFP-HvBDG1 (Fig 2e,f). However, since Ara6 and Ara7 only label a subset of endomembrane compartments, we cannot exclude that RFP-HvBDG1 localises to a different set of endosomes not shared with Ara6 and Ara7 (Fig 2g,h). Overall, these data suggest that HvBDG1 does not follow the conventional secretory pathway through the trans-Golgi network but may travel through an unconventional secretory pathway associated with the ER and mobile bodies which travel along the ER tubular network. We did not detect signal in the outer cell wall as reported for Arabidopsis

**Figure 2.**
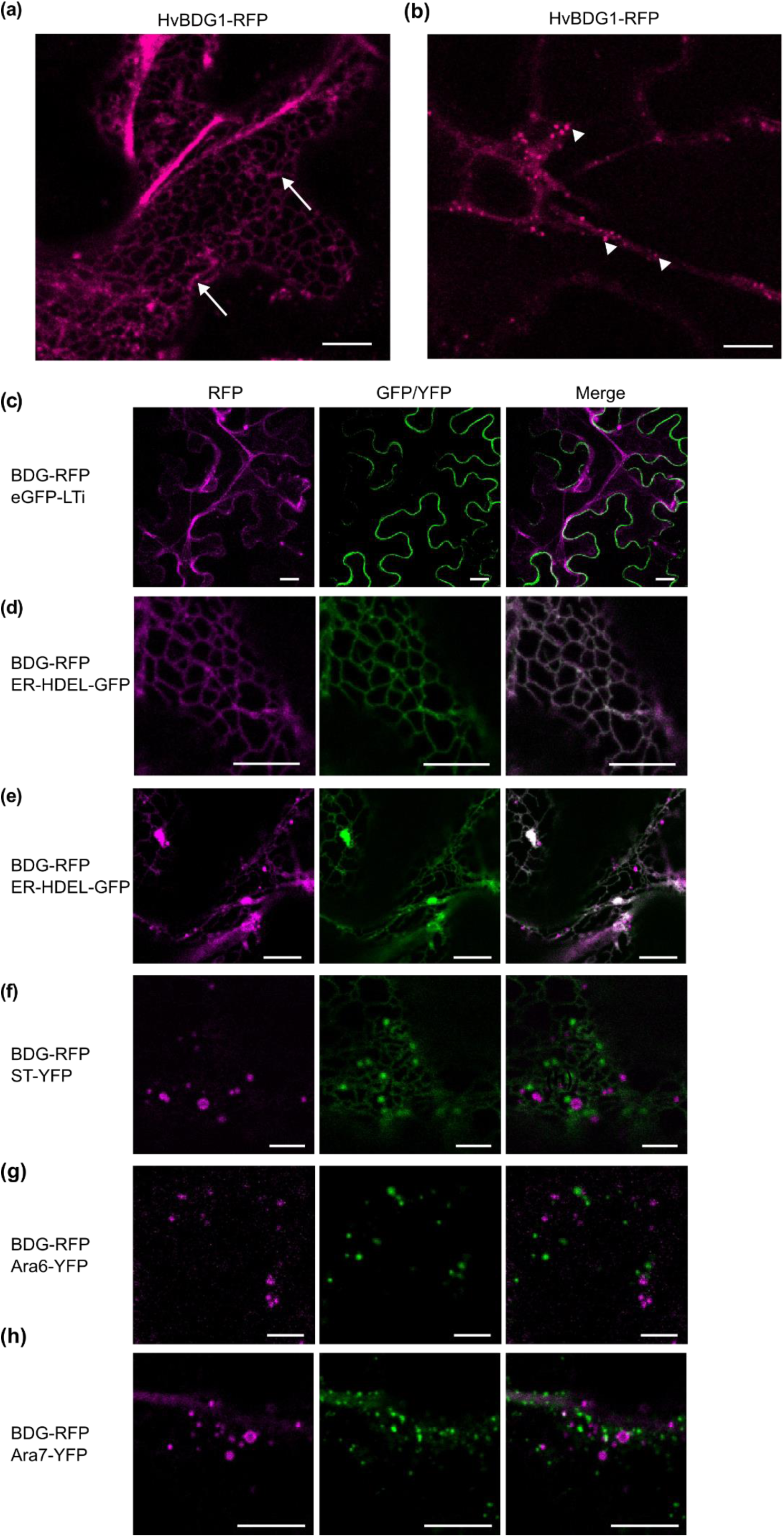
Tagged HvBDG1 protein localises to the endoplasmic reticulum and spherical mobile bodies. Localisation in transiently infiltrated *Nicotiana benthamiana* leaves expressing constructs as indicated. (a) shows leaves 2 days post-infiltration (DPI); all other panels show samples are 3 DPI. (a-b) Red Fluorescent Protein (RFP)-HvBDG1. Reticulate structures characteristic of the endoplasmic reticulum (ER) indicated by arrows and spherical bodies by arrowheads. Scale bars, 10 μm. (c) RFP-HvBDG1 expressed in leaves of a stable *N*. *benthamiana* line expressing Lti-PM-GFP marker for the plasma membrane. Scale bars, 20 μm. (d) RFP-HvBDG1 expressed in leaves of a stable *N*. *benthamiana* line expressing green fluorescent protein (GFP) fused to an ER retention motif (ER-HDEL-GFP). Scale bars, 10 μm. (d) (e) RFP-HvBDG1 expressed in stable *N*. *benthamiana* line expressing ER-GFP showing association of spherical mobile bodies (arrows). (f) RFP-HvBDG1 and the Golgi marker ST-YFP. Scale bars, 5 μm. (g) RFP-HvBDG1 and the endosomal marker Ara6. Scale bars, 5 μm. (h) RFP-HvBDG1 and the endosomal marker Ara7. Scale bars, 10 μm.

### BDG1 and WIN1 promotion of the wax bloom is conserved in wheat

We tested whether *BDG* and *WIN* genes similarly promote wax blooms in other cereal species by developing durum wheat mutant lines. While barley is diploid, durum is an allotetraploid species consisting of A and B sub-genomes. For *HvBDG1* (located on the short arm of chromosome 7H at 18.1 Mb), collinearity between barley and wheat (e.g. Wang et al., 2015) expects the orthologous durum wheat homoeologues to be located on the short arms of chromosomes 7A and 7B. Analysis of the durum reference genome found that while *TdBDG1-A1* (*TRITD7Av1G017740*) is located at the expected colinear location (chromosome 7A at 32.1 Mb), the expected 7B homoeologue actually resided on the long arm of chromosome 4A at 685.5 Mb (*TRITD4Av1G244120*, termed here *TdBDG1-A2*), due to the chromosomal translocation T(4AL;7BS)1 present in both tetraploid (Dvorak et al., 2018) and hexaploid (Zhou et al., 2020) wheat (Supplemental Note 2). Thus, from an evolutionary perspective, *TdBDG1-A2* represents the durum wheat B sub-genome homoeologue, rather than a *BDG1* paralogue. For *HvWIN1* (chromosome 6H at 187.9 Mb), durum orthologues were identified at the expected colinear locations on the short arms of chromosomes 6A (*TRITD6Av1G082070* at 204.1 Mb; *TdWIN1-A*) and 6B (*TRITD6Bv1G086320* at 262.3 Mb Mb; *TdWIN1-B*). Next, TILLING mutants containing disruptive mutations in the durum homoeologues were identified: for *TdBDG1*, two independent *TdBDG1-A1* premature stop mutants (*TdBDG1-A1^STOP75^* caused by a nonsense G225/A mutation in the CDS within exon-1, resulting in a stop codon at amino acid residue 75 in the predicted protein, W75/*, TILLING line Kronos3179. *TdBDG1-A1^STOP385^* CDS position G1154/A, W385/*, Kronos0599) and one premature stop mutant in *TdBDG1-A2* (*TdBDG1-A2^SPLICEi2d^,* due to a GT/AT mutation in the intron-2 canonical splice donor motif, resulting in a nonsense mutation in the subsequent codon that terminates the predicted protein at amino acid residue 320. Kronos2087) (Fig 3a). All three mutants resulted in either complete (*TdBDG1-A1^STOP75^, TdBDG1-A1^STOP385^*) or partial (*TdBDG1-A2^SPLICEi2d^*) loss of the alpha/beta hydrolase fold-1 predicted protein domain (pfam ID PF00561) (Fig 3a). For *TdWIN1,* three TILLING lines were selected, all of which contained mutations in the AP2/ERF predicted protein domain (Panther ID PTHR31194): premature stop mutant *TdWIN1- A^STOP125^*(Kronos3416) and two missense mutations with SIFT scores = 0 at conserved amino acid residues on the A (*TdWIN1-A^G130/E^*, Kronos2646) and B (*TdWIN1-B^P26/A^*, Kronos2314) homoeologues (Fig 3b). For each target durum gene, mutated homoeologues were combined into single genetic backgrounds by crossing to generate F_1_ hybrids, followed by marker-assisted selection in F_2_ progeny derived from selfed F_1_ plants. For *TdBDG1*, homozygous *TdBDG1-A1 and TdBDG1-A2* double mutants (combinations *TdBDG1-A1^STOP75^/TdBDG1-A2^SPLICEi2d^* and *TdBDG1-A1^STOP385^/TdBDG1-A2^SPLICEi2d^*) all lacked the wax blooms on the spike, stem and leaf sheath observed in the wild-type segregants (Fig 3c; Fig S8). All *TdWIN1* double mutants lacked wax blooms on the ear and stem (Fig 3d; Fig S8). Accordingly, analysis of the durum mutants confirmed conservation of *BDG1* and *WIN1* function between barley and wheat. The ability to generate a more quantitative effect on durum wax phenotype was shown via comparison of the different mutant combinations. For example, in single *TdBDG1-A1* homoeologue mutants, loss of wax phenotype in the spike was more pronounced in the mutant carrying a stop mutation closer to the predicted protein N-terminus (*TdBDG1-A1^STOP75^*) compared to the *TdBDG1- A1^STOP385^* mutant that retained a larger proportion of the protein. While the *TdBDG1-A2 ^SPLICEi2d^* mutant in the second durum homoeologue did not show a notable loss of cuticular wax in the tissues observed, mutation in both homoeologues was nevertheless necessary to obtain a waxless phenotype (Fig 3c). Similar quantitative effects on wax phenotype were also observed for different combinations of *TdWIN1* mutants, especially in the ear and stem (Fig 3d), with missense/missense combination *TdWIN1-A^G130/E^*/*TdWIN1-B^P26/A^* notable for retaining reduced leaf sheath surface wax deposition compared to complete loss of leaf sheath wax in *TdWIN1-A^STOP125^*/*TdWIN1-B^P26/A^*(Fig S8). We also confirmed that leaf blades from both *TdBDG1* double mutants showed increased permeability to Toluidine Blue, suggesting that cuticle integrity is defective in these lines (Fig S9).

**Figure 3.**
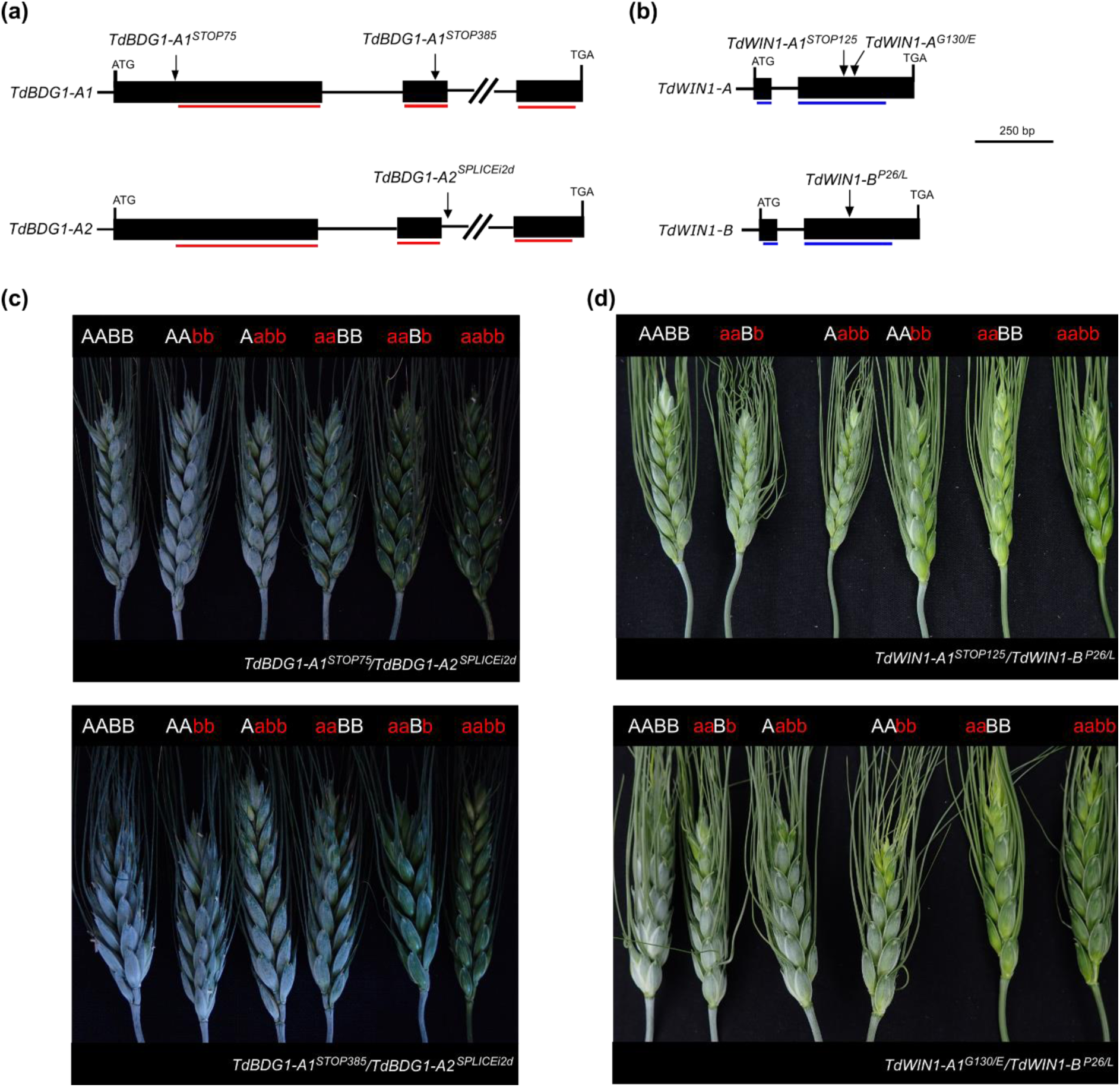
Durum wheat *TdBDG1* and *TdWIN1* TILLING mutants confirm conserved role in the control of epidermal wax bloom between wheat and barley. Exon/intron structure of durum wheat A- and B-subgenome homoeologues of (a) *TdBDG1* and (b) *TdWIN1*. The locations of the selected TILLING mutants are indicated. The exon regions that code for predicted protein domains in *TdBDG1* (alpha/beta hydrolase fold-1 domain, pfam ID PF00561) and *TdWIN1* (AP2/ERF domain, Panther ID PTHR31194) are shown using red and blue horizontal bars, respectively. Examples of wax bloom phenotype observed in durum wheat spikes from F_2_ individuals carrying different wild-type and mutant alleles at homoeologues of (c) *TdBDG1* and (d) *TdWIN1*. Two combinations of mutants are shown for both *TdBDG1* and *TdWIN1* - one in the upper panels and a second in the lower panels. For each combination, allelic state at the A- and B- sub-genome homoeologue is indicated via uppercase (wild-type) or lowercase (mutant) letters, such that homozygous wild-type (AA or BB), homozygous mutant (aa or bb), and combinations of homozygous and wild type (e.g., AABb) alleles are indicated.

### HvBDG1 or HvWIN1 prevent grain skinning in barley

In addition to glossy and permeable cuticles, *hvbdg1^156^* also showed grain skinning (Fig 4a; Fig S10), a distinctive cuticular specialisation not found in wheat. Manual threshing revealed that 49% and 47% of *hvbdg1^156^* and *hvbdg1^406^* grain, respectively, shed hulls compared to only 7% of Bowman (Fig 4b). Following separation in a debranner (assay described in Campoli et al., 2024), *hvbdg1^156^* and *hvbdg1^406^* grain lost 85% and 86% of its hulls by weight, respectively, compared to 73% hulls shed from Bowman grain (p < 0.001; Fig 4c). We noticed that *hvwin1^126^* and *hvwin1^407^* also skinned (Fig S10) with 46% of *hvwin1^126^* grained skinning upon manual threshing (Fig 4b), suggesting that *hvwin1* alleles also share this *hvbdg1* phenotype. Grain from *hvwin1^126^* and *hvwin1^407^* were statistically equivalent or showed more hull loss compared to Bowman in the debranner depending on the experiment (Fig 4b). Together, these results indicate HvBDG1 and HvWIN1 are essential for strong hull to caryopsis adhesion.

**Figure 4.**
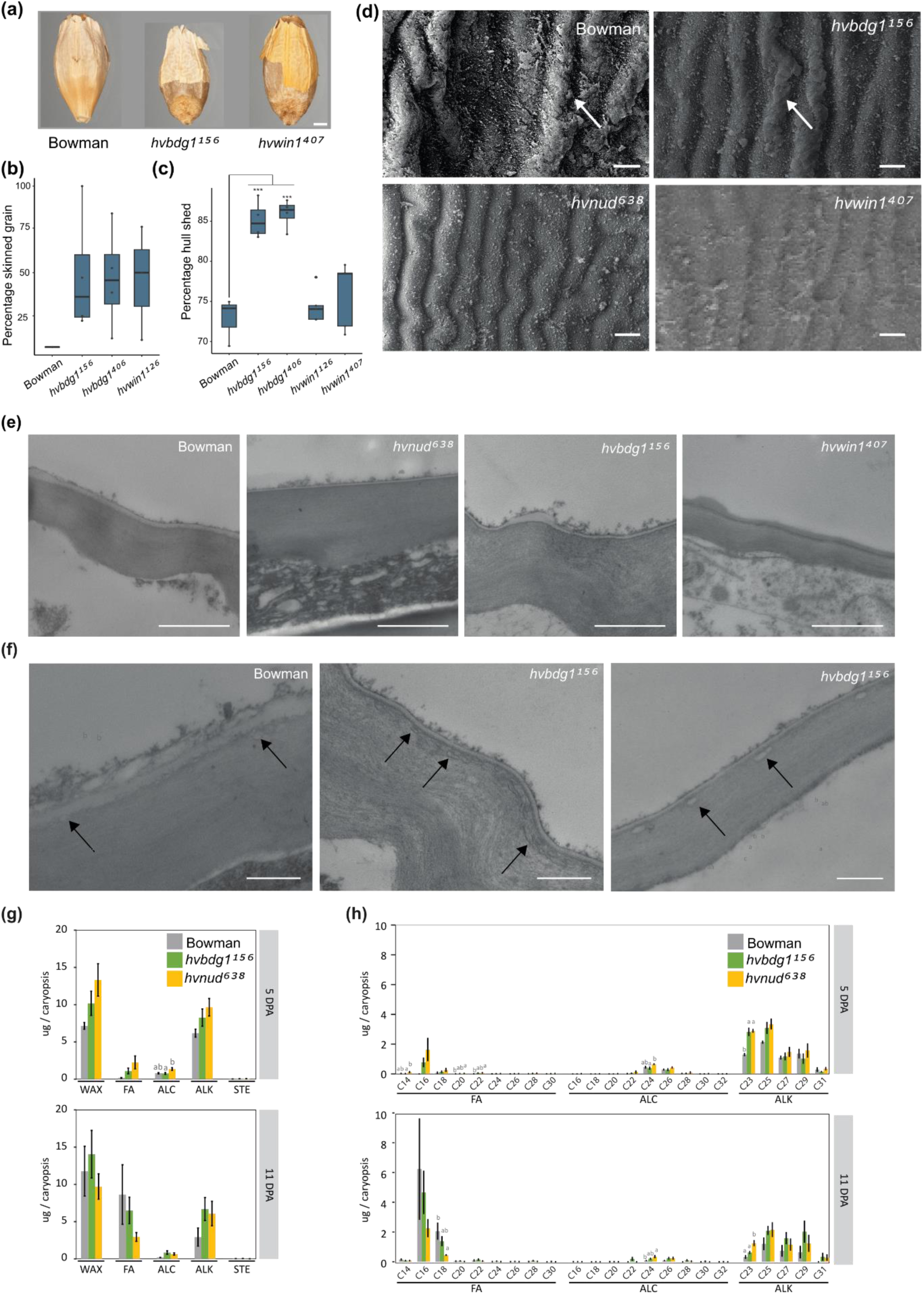
*HvBDG1*, *HvWIN1* and *NUD* regulate grain skinning and features of the caryopsis pericarp cuticle. (a) Photos of hull adhesion in Bowman, *hvbdg1^156^* and *hvwin1^406^* manually threshed grain. Dorsal side. Scale bars, 0.05 cm. (b) Manual assay indicates percentage of grain showing any skinning. Bowman (n = 1 from multiple plants), *hvbdg1^156^* (n = 4), *hvbdg^406^* (n = 4) and *hvwin1^126^* (n = 3), *hvwin1^407^* (n=5). (c) Debranner assay showing extent of hull separation in Bowman (n = 3), *hvbdg1^156^* (n = 4), *hvbdg^406^* (n = 4) and *hvwin1^126^* (n = 4), *hvwin1^407^* (n = 5). Boxes represent the interquartile range with the horizontal line representing the median. Whiskers represent the upper and lower quartiles plus 1.5 times the interquartile range. Points represent individual replicates. *** p < 0.001 (ANOVA followed by Tukey’s HSD Post-Hoc test). (d) Scanning electron microscopy (SEM) of 11 days post anthesis (DPA) caryopsis (hulls removed) pericarp surfaces in Bowman, *nud^638^*, *hvbdg1^156^* and *hvwin1^407^*. Arrows indicate accumulating cuticular material on ridges. Scale bars, 1 μm. (e) Transmission electron microscopy (TEM) of the outer pericarp in sections made from 11 DPA caryopses (hulls removed) in Bowman, *nud^638^*, *hvbdg1^156^* and *hvwin1^407^*. Scale bars, 1 μm. (f) Higher magnification TEM of the pericarp cell wall-cuticle interface of 11 DPA caryopses (hulls removed) in Bowman and *hvbdg1^156^*. Scale bars, 500 nm. Images are representative of at least three biological replicates. (g) Soluble wax classes and (h) Soluble wax chain lengths detected in 5 and 11 DPA caryopses from Bowman, *nud^638^* and *hvbdg1^156^*. WAX = total wax extracted, FA = fatty acids, ALC = primary alcohols, ALK = alkanes, STE = sterols. Bars show the average with standard deviation of three biological replicates. Letters indicate significant differences within genotypes (P < 0.05; Tukey’s HSD multiple comparison following one-way ANOVA).

### *HvBDG1*, *HvWIN1* and *NUD* regulate pericarp surface features during hull adhesion

Deletion of *NUD* leads to a complete loss of hull attachment while impaired HvWIN1 and HvBDG1 function weakens hull adhesion. We hypothesised that *HvWIN1* and *HvBDG1* could be part of a NUD regulatory pathway to modify the pericarp cuticle, where loss of function in either leads to a partially defective pathway and skinning. We predicted that a partially defective pathway may lead to skinning mutants with pericarp features intermediate between naked and covered grain. To test this hypothesis, we used scanning electron microscopy (SEM) to examine the outer pericarp surfaces of caryopses from a near isogenic line panel of *hvbdg1^156^* and *hvwin1^407^* skinning alleles, the naked BW- NIL638 (*nud^638^*) allele and Bowman. We selected caryopses at 7 DPA, prior to hull adhesion and at 11 DPA during hull adhesion (hulls show first signs of adhesion around 9 DPA). At 7 DPA, Bowman pericarp epidermal cells showed a mixture of smooth surfaces, narrow perpendicular nanoridges and wider longitudinal ridges; *hvwin1^407^* resembled Bowman while *hvbdg1^156^* had some instances of crooked longitudinal ridges and *nud^638^* had cells showing perpendicular striations (Fig S11). At 11 DPA, waxy plaques decorated the Bowman pericarp cell ridges but were infrequent and smaller in *hvbdg1^156^* and *hvwin1^407^*, and not readily observed in *nud^638^* (Fig 4d). Transmission electron microscopy (TEM) on 7 DPA Bowman caryopses showed thick cuticles with underlying and darkly stained electron-lucent globules and fibrillar extrusions that extended from the cell wall of Bowman which were not present in BW638 which instead had a thin cuticle (Fig S12). By 11 DPA, Bowman had very thick pericarp cuticles with outer lamellations and an internal region of merged cell-wall cuticle interface containing electron-lucent globules (Fig 4e,f). In contrast, pericarp cuticles in 11 DPA *nud^638^* samples were thin with a sharp and smooth electron dense boundary to the outer cell wall lacking globules or fibrillar extensions (Fig 4e). Pericarp cuticles in *hvbdg1^156^*, and to a lesser extent *hvwin1^407^*, were thin but with occasional thickened regions and smooth, electron dense cell wall-cuticle interfaces (Fig 4e). The *hvbdg1^156^* pericarp cell wall showed electron-lucent globules which did not merge with the cuticle, in contrast to Bowman (Fig 4e,f). Taken together, these data indicate that NUD, HvBDG1 and HvWIN1 regulate pericarp changes coincident with hull adhesion including cuticular thickening, appearance of epicuticular plaques and integration between the cuticle and the outer cell wall. Consistent with our hypothesis, these features were more disrupted in *nud^638^* compared to *hvbdg1^156^* and *hvwin1^407^* which showed intermediate phenotypes compared to Bowman.

### Roles of *HvBDG1*, *HvWIN1* and *NUD* in pericarp and hull surface composition

We next asked whether changes in pericarp surface and ultrastructure correlated with changes in cuticle composition. We used Gas Chromatography-Mass Spectrometry (GC-MS) to compare soluble surface lipids and cutin monomers extracted from 5 and 11 DPA caryopses (hulls removed) and 11 DPA hulls from our NIL panel. We examined Bowman, *nud^638^* and *hvbdg1^156^* in one experiment and Bowman and *hvwin1^407^* in another experiment, therefore these data are shown on separate graphs (Fig 4g,h; Table S6; Fig S13). Data from Bowman and *nud^638^* were published in a previous report (Campoli et al., 2024) and are used here for comparative analyses. In all genotypes, alkanes dominated soluble lipids extracted from caryopses at 5 DPA, followed by fatty acids and alcohols, with only small amounts of sterols. We detected differences in specific chain lengths with increased shorter length alkanes and fatty acids in naked and skinning lines. For instance, *nud^638^* (117%, p < 0.001), *hvbdg1^156^* (122%, p < 0.001) and *hvwin1^407^*(58%, p = 0.049) accumulated more C23 alkanes at 5 DPA compared to Bowman (Fig 4h; Fig S9). Although this moiety decreased in all genotypes by 11 DPA, C23 alkanes remained higher in *nud^638^* (279%, *p* = 0.003) and *hvwin1^407^*(113%, p = 0.018) compared to Bowman (Fig 4f; Fig S13). C14 fatty acids were increased specifically in *nud^638^* at 5DPA (333%, p = 0.026), and while C16 fatty acids accumulated highly in both *hvbdg1^156^* and *nud^638^*, they remained undetectable in Bowman. We compared 5 DPA and 11 DPA for changes coincident with hull adhesion (Table S6). Total soluble lipids showed no change; however, Bowman, *hvbdg1^156^* and *hvwin1^407^* samples showed increased total fatty acids from 5 DPA to 11 DPA, while this change was not observed in *nud*^638^ (Table S6). Alcohols, in particular C24, decreased in Bowman, *nud^638^* and increased in *hvwin1^407^* between 5 and 11 DPA and resulted in an increased amount in *nud^638^* compared to Bowman at 11DPA (338%, p = 0.012; Table S6, Fig 4h). While found in small amounts, sitosterol also increased by 215% in *nud^638^* compared to Bowman (p = 0.037; Fig S13). We also measured cutin components (Fig S14; Table S6). Consistent with our previous results (Campoli et al., 2024), the major components were mono and di- hydroxy C16 and C18 acids, with a prevalence of C16 moiety, followed by the C18 mono and di-hydroxy acids (Fig S14; Table S6). Other than a significantly higher amount of ωOH C16 (ω- hydroxyhexadecanoic acid) in *hvbdg1^156^* at 5 DPA, all lines showed a similar composition and increased these components between 5 and 11 DPA (Fig S14; Table S6). Altogether, skinning and naked lines showed overlapping as well as genotype-specific statistical shifts towards shorter chain alkanes and fatty acids from 5 to 11 DPA, but little change in cutin components, suggesting little association between cutin and hull adhesion.

As expected from their glossy phenotypes, hull extracts had less total wax content in *hvbdg1^156^* and *hvwin1^407^* compared with Bowman, mostly due to loss of β-diketones and hydroxy β-diketones (Fig S13). Compared to Bowman, alcohols, alkanes and fatty acids in *hvbdg1^156^* and *hvwin1^407^* shifted from longer chain towards shorter chain lengths, a trend also detected in *nud^638^*, despite no reported phenotype for *nud^638^* outside the grain (Fig S13). Major cutin components detected in hulls at 11 DPA were aromatics, in particular ferulic and coumaric acid, followed by ωOH C16. As for caryopses, cutin composition in hulls did not differ significantly in the mutants compared to Bowman (Fig S14). Taken together, NUD, HvBDG1 and HvWIN1 share roles in extending longer chain aliphatics in both hull and caryopses tissues, in particular alkanes and fatty acids, while HvBDG1 and HvWIN1 are essential to promote β-diketone deposition on hulls.

### *HvWIN1* and *NUD* regulate overlapping and distinct genes during grain development

AtWIN/SHN transcription factors upregulate *BDG* expression in Arabidopsis (Shi et al., 2011), so we hypothesised that NUD and/or HvWIN1 similarly promote *HvBDG1* expression. We first examined *HvBDG1* expression by qRT-PCR over a developmental time course in Bowman caryopses from 3-10 DPA. *HvBDG1* transcripts peaked coincident with the first signs of hull adhesion at 9 DPA (Fig 5a). We then selected two time points to compare *HvBDG1* expression in *nud^638^* and *hvwin1^407^*. At 5 DPA, *nud^638^* and *hvwin1^407^* caryopses had 70% and 60% *HvBDG1* expression compared to Bowman, respectively (Fig 5b), while at 9 DPA, *HvBDG1* expression declined 50% in *nud^638^*, with little change in *hvwin1^407^* (Fig 5c), supporting that both HvWIN1 and NUD influence *HvBDG1* expression early in grain development with a major role for NUD as hulls adhere.

**Figure 5.**
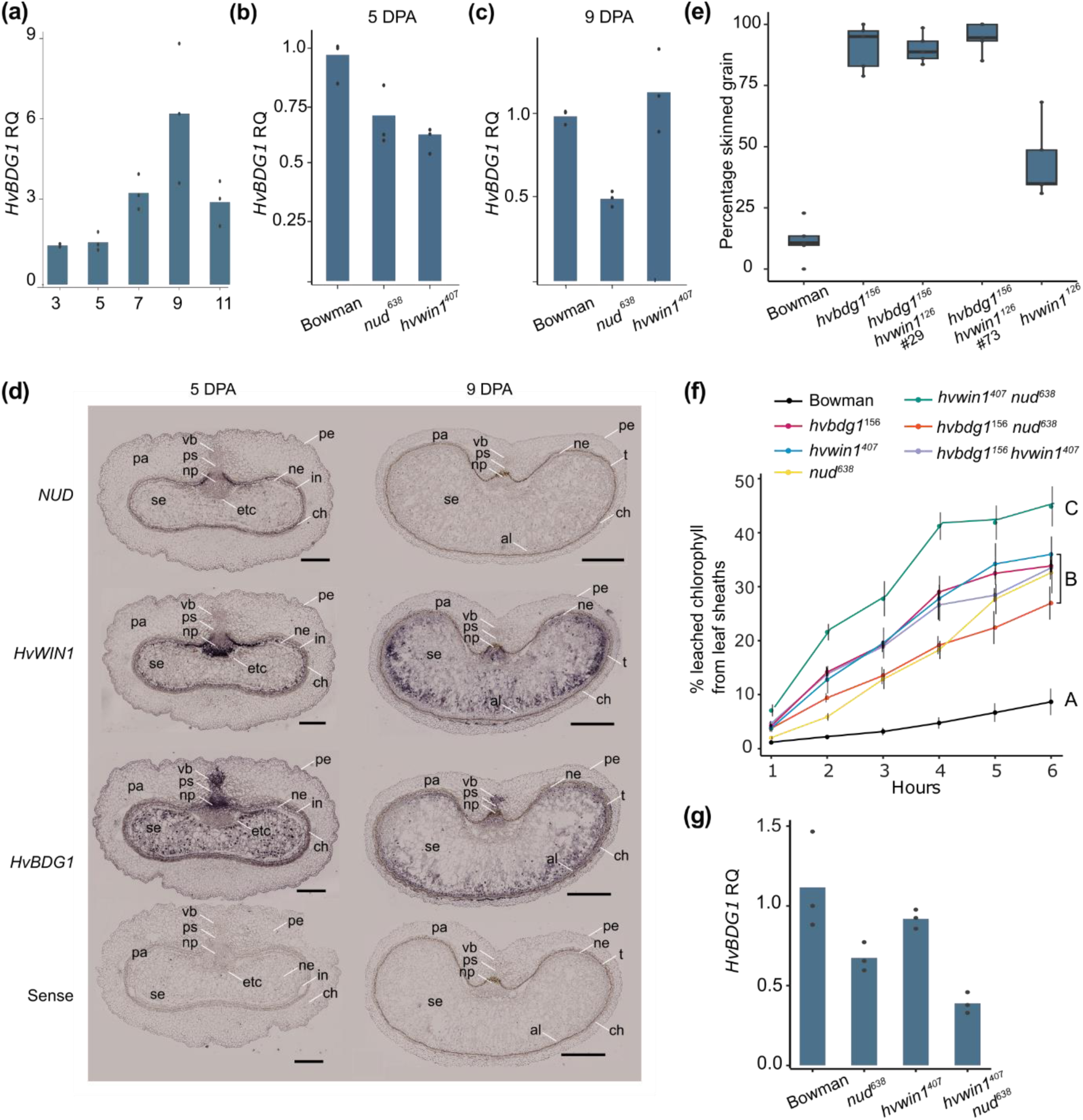
*HvBDG1*, *HvWIN1* and *NUD* expression and interaction. (a-c) qRT-PCR of *HvBDG1* transcripts in: (a) developmental time course of Bowman caryopses 3 to 11 days post anthesis (DPA); (b) 5 DPA in Bowman, *hvwin1^407^* and *nud^638^* caryopses, and (c) 9 DPA in Bowman, *hvwin1^407^* and *nud^638^* caryopses. (d) RNA *in situ* hybridisation of caryopses sectioned from Bowman at 5 and 9 DPA. Sections hybridised with antisense probes specific for *NUD*, *HvWIN1* and *HvBDG1* transcripts, or a non-specific sense probe as indicated. Labels are pe, pericarp epidermis; pa, pericarp parenchyma; ch, pericarp chlorenchyma; ps, pigment strand; np, nucellar projection; n, nucellus; t, testa; etc, endosperm transfer cells; al, aleurone. Scale bars, 200 μm (5 DPA) and 500 μm (9 DPA). (e) Skinning index of Bowman, single (*hvbdg1^156^*, *hvwin1^407^*) and double mutants (*hvbdg1^156^ hvwin1^407^*). (f) Chlorophyll leaching assays of leaf sheaths from Bowman, single mutants (*hvbdg1^156^*, *hvwin1^407^*, *nud^638^*), and double mutants (*nud^638^ hvwin1^407^*, *hvbdg1^156^ nud^638^* and *hvbdg1^156^ hvwin1^407^*). (g) qRT-PCR of *HvBDG1* transcripts in Bowman, *hvwin1^407^*, *nud^638^* and *nud^638^ hvwin1^407^* base tissue from the second leaf blade.

Since HvWIN1 and NUD influence *HvBDG1* expression in grain, and all three genes control hull adhesion, we next asked whether their spatial expression overlaps in 5 DPA and 9 DPA caryopses using RNA *in situ* hybridisation. At 5 DPA, *NUD* transcripts were detected in maternal tissues including the integuments and the nucellar epidermis, most strongly next to the nucellar projection, as well as faintly in the outer pericarp, vascular bundle and pigment strand (Fig 5d). *HvWIN1* and *HvBDG1* transcripts also accumulated in these tissues at 5 DPA, with additional strong signals for *HvWIN1* in the lower and *HvBDG1* in the upper nucellar projection as well as in the pigment strand and vascular bundle (Fig 5d). By 9 DPA, the nucellar epidermis compresses against the testa formed through integument maturation. We detected no signal in the testa at 9 DPA for any probe, and we speculate that the characteristic pigment accumulation during maturation may interfere with probe hybridisation and/or detection. At 9 DPA, *HvWIN1* and *HvBDG1* transcripts accumulated in endosperm and the nucellar projection with *HvBDG1* still detected in the pigment strand as well as the aleurone; interestingly *NUD* was not detected at this stage (Fig 4d). Taken together, *HvWIN1*, *NUD* and *HvBDG1* expression overlaps in the integuments, nucellar epidermis and the pericarp epidermis, while *HvWIN1* and *HvBDG1* share expression in the endosperm and the nucellar projection.

### *HvWIN1* and *NUD* independently control cuticle properties

To learn more about possible shared functions between *HvBDG1*, *HvWIN1* and *NUD* and their interactions, we used genetic analyses by generating double mutants from the NILs. We observed that the *hvbdg1^156^ hvwin1^407^* double mutant skinned equivalently to *hvbdg1^156^* while *hvwin1^407^* skinned intermediate to *hvbdg1^156^* and Bowman grain (p < 0.001, Tukey HSD; Fig 5e; Fig S10), supporting the hypothesis that HvBDG1 may work downstream of HvWIN1 in the control of hull adhesion, and that other factors besides HvWIN1 contribute to *HvBDG1* expression in *hvwin1^407^* grain. Combining *hvwin1^407^* or *hvbdg1^156^* with *nud^638^* broadly reflected in an additive effect on phenotype, with *hvbdg1^156^nud^638^* and *nud^638^ hvwin1^407^* double mutant grain always naked and *hvbdg1^156^ nud^638^* and *nud^638^ hvwin1^407^* leaf sheaths equivalently glossy to the *hvbdg1^156^* or *hvwin1^407^* glossy parent. Measuring leaf cuticle permeability in Bowman, single mutants and double mutants revealed increased permeability in *nud^638^* leaf sheaths, potentially showcasing a previously unknown role for NUD outside of the grain in the promotion of leaf cuticle integrity (Fig 5f). Moreover, *nud^638^ hvwin1^407^* double mutant leaf sheaths and leaf blades were more permeable than either parent (*p* = 0.000 and *p* = 0.038, Fig 5b; Fig S15), indicating that WIN/SHN factors in barley non-redundantly control cuticle integrity throughout the leaf. We did not see any compensatory increase in *HvWIN1* levels in *nud^638^* leaves or vice versa, suggesting that NUD and HvWIN1 do not regulate each other’s expression in leaves (Fig S16). Combining either *nud^638^* or *hvwin1^407^* with *hvbdg1^156^* increased leaf blade permeability compared to either parent (Fig 5f), suggesting additive roles, while leaf sheaths of the double mutants *hvbdg1^156^ nud^638^* and *hvbdg1^156^ hvwin1^407^* showed the same increased permeability as parent lines compared to wild type, potentially indicating that HvBDG1 works in the same pathway as the WIN/SHN factors in the leaf sheath. We compared *HvBDG1* expression in second leaf blades separated into a basal proximal section containing dividing and younger differentiating cells and a middle section with differentiating cells thickening their cuticle. In basal sections, *HvBDG1* transcripts accumulated to 80% and 70% of Bowman levels in *hvwin1^407^* and *nud^638^*, respectively (Fig 5g), while the leaf blade mid-section accumulated *HvBDG1* mRNA to 60% of Bowman levels in *hvwin1^407^* with no change in *nud^638^*(Fig S17). We observed more severe suppression of *HvBDG1* expression in the *nud^638^ hvwin1^407^* double mutant leaf bases (Fig 5g), supporting a correlation of WIN/SHN promotion of *HvBDG1* expression and cuticular integrity. Taken together, double mutant analyses revealed non-redundant roles for HvWIN1 and NUD in leaf cuticular integrity and suggested that HvBDG1 may work downstream of these factors to control leaf integrity and hull to caryopsis adhesion, supported by comparative qRT-PCR indicating that both HvWIN1 and NUD regulate *HvBDG1* expression, with HvWIN1 more important in leaves and NUD more important in caryopses.

### HvWIN1 and NUD control overlapping and distinct transcriptomes in developing grain

*HvWIN1* and *NUD* have overlapping but divergent expression patterns in the developing grain and likely regulate many genes besides *HvBDG1*. To learn more about gene expression sensitive to NUD and HvWIN1 function, we performed RNA-seq on Bowman, *nud^638^* and *hvwin1^407^* caryopses at 5 DPA and 9 DPA. Compared to Bowman, we detected 652 and 575 differentially expressed genes (DEGs) at 5 DPA, and 424 and 690 DEGs at 9 DPA in *nud^638^* and *hvwin1^407^*, respectively (Fig 6a; Table S7). We examined all DEG lists for overlap between stages and genotypes, producing a Venn diagram showing these relationships (Fig 5a; Table S7). Of the 26 shared DEGs across all stages and genotypes, we observed common downregulation of barley *PROTODERMAL FACTOR 1* (*HvPDF1*; *HORVU.MOREX.r3.3HG0237080*), a marker for specification of epidermal tissues and cuticle formation in Arabidopsis (Abe et al., 1999), as well as severe downregulation of *TRICHOME BIREFRINGENCE-LIKE 19* (*TBL19*) involved in cellulose deposition and pectin cross-linking with the cell wall (*HORVU.MOREX.r3.2HG0167810*, Bischoff et al., 2010); highly upregulated genes included two encoding NBS-LRR disease resistance proteins RPM1 (*HORVU.MOREX.r3.2HG0199120* and *HORVU.MOREX.r3.7HG0735600*) and a stress-related cellulase (*HORVU.MOREX.r3.1HG0000950*), together suggesting changes in gene expression consistent with epidermal differentiation and cuticle/cell wall damage (Fig 5b). We also examined DEGs shared in three of four categories. Here we observed differential expression of cell wall-associated genes, lipid metabolism and transport, disease resistance and wound response genes as well as signalling genes (Fig 6b). The 180 DEGs shared between *nud^638^* and *hvwin1^407^* at 5 DPA (Fig 6c) included further downregulation of genes encoding multiple GDSLs and proline-rich proteins normally localised to the cell wall, and upregulation of a fatty acyl-CoA reductase catalysing primary alcohol production in Arabidopsis seed coats (FAR4, *HORVU.MOREX.r3.4HG0414900*; Vishwanath et al., 2013) which may contribute to increased accumulation of C24 alcohols in *nud^638^* and *hvwin1^407^* 11 DPA caryopses (Table S6). We also detected increased transcript levels for a gene encoding a GASSHO-Like receptor kinase (*HORVU.MOREX.r3.7HG0704310*) whose homologues are in cuticle maintenance within Arabidopsis seeds (Doll et al., 2020). At 9 DPA, shared DEGs included upregulated genes encoding putative lipid transfer proteins (LTPs) and beta-glucosidase genes involved in cell wall metabolism, as well as downregulation of another *TBL19* homologue (Fig 5d). (Fig 5d). Downregulated DEGs specific to *nud^638^* at either stage (Fig 6f) included cuticular machinery genes encoding β-ketoacyl-CoA synthases (KCSs) catalysing the first step in fatty acyl elongation: HvKCS1 (*HORVU.MOREX.r3.4HG0392320*) whose defective alleles underlie *cer-zh* (Li et al., 2018), homologues of *KCS5* (*HORVU.MOREX.r3.6HG0612530*) and *KCS6* (*HORVU.MOREX.r3.7HG0670360*), as well as the barley *β-ketoacyl-CoA reductase 1* homolog (*KCR1*; *HORVU.MOREX.r3.7HG0653780*), consistent with fewer fatty acids in *nud^638^*. We also detected downregulation of *HvABCG31* (*HORVU.MOREX.r3.3HG0240110*), encoding a putative transporter responsible for the reduced cutin barley mutant *eibi1*, the gene *HvCER3.2* (*HORVU.MOREX.r3.7HG0741770*; Chen et al., 2011), whose Arabidopsis homologue encodes part of a VLC alkane synthase complex (Bernard et al., 2012), as well as genes for a glycosylphosphatidylinositol (GPI)-anchored lipid transfer protein, *LTPG2* (*HORVU.MOREX.r3.3HG0296370*), linked to cuticular wax export/accumulation in Arabidopsis (Kim *et al*., 2012) and a putative barley *CUTIN SYNTHASE* (*CUS*; *HORVU.MOREX.r3.5HG0486520*) whose homologues polymerise cutin monomers in Arabidopsis and tomato (Yeats et al., 2012; Hong et al., 2017; Sagado et al., 2020). Upregulated genes in *nud^638^* include homologues of *HOTHEAD* paralogue (*HORVU.MOREX.r3.5HG0475790*), a gene associated with cutin synthesis in Arabidopsis and rice (Kurdyukov et al., 2006b; Xu et al., 2017), and the *HOTHEAD* paralog *IPE1* linked to cutin synthesis in maize (*HORVU.MOREX.r3.2HG0186860*; Chen *et al*., 2017), two further barley *ABCG11/12* genes (*HORVU.MOREX.r3.2HG0183620*, *HORVU.MOREX.r3.1HG0032470*), as well as homologues encoding epidermal and integument developmental regulators, such as GLABROUS11 (HDG11; *HORVU.MOREX.r3.1HG0006070*; Khosla et al., 2014) and the MYB transcription factor KANADI4 (*HORVU.MOREX.r3.5HG0513030*; McAbee et al., 2006). Impaired HvWIN1 function causes misexpression of a similar but distinct suite of cuticular genes and other developmental regulators (Fig 6g), including downregulation of a barley *glycerol-3-phosphate acyltransferase6* (*GPAT6*; *HORVU.MOREX.r3.4HG0340350*) related to genes important for cutin accumulation and cell wall properties in tomato (Petit et al., 2016; Fawke et al., 2019) and cutin dependent nanoridge formation in Arabidopsis (Li-Beisson et al., 2009), and abolished expression of a *phosphoethanolamine N-methyltransferase* (*NMT1*; *HORVU.MOREX.r3.6HG0573780*) associated with endomembrane trafficking in Arabidopsis (Renna et al., 2013). We also detected upregulation of *KCS17* (*HORVU.MOREX.r3.5HG0474980*) important for waxes in Arabidopsis seed coats (Kim et al., 2024), two wax synthases (*HORVU.MOREX.r3.2HG0138620* and *HORVU.MOREX.r3.7HG0665520*), another homologue of *PDF1* (*HORVU.MOREX.r3.3HG0286190*), *HDG11* (*HORVU.MOREX.r3.6HG0618540*) and *KANADI2* (*HORVU.MOREX.r3.6HG0606010*; McAbee et al., 2006), as well as downregulation of *BEL1* (*HORVU.MOREX.r3.1HG0052320*; Reiser et al., 1995). The *HvWIN1* gene itself was also upregulated, suggesting autoregulation (Table S7). Compared to Bowman, *HvBDG1* expression was 70% lower in both *hvwin1^407^* and *nud^638^* at 5 DPA (*p* = 7.68 x10^-6^ and *p* = 2.24 x10^-4^) and reduced 80% and 50% in *hvwin1^407^* and *nud^638^* at 9 DPA (p = 0.030 and p = 2.99 x10^-5^, respectively; Table S7), broadly in agreement with qRT-PCR (Fig 5). RNA-seq also showed that *NUD* levels do not increase in *hvwin1^407^* but a modest 1.4 increase in *HvWIN1* in *nud^638^* at 9 DPA (*p* = 0.005; Table S7), suggesting that *HvWIN1* expression may be induced with loss of *NUD* and that function of NUD is not required for *HvWIN1* induction or vice versa. In sum, comparative RNA-seq suggested that NUD and HvWIN1 function regulates shared and distinct target genes, including *HvBDG1*, many with functions associated with cell wall and cuticle formation.

**Figure 6.**
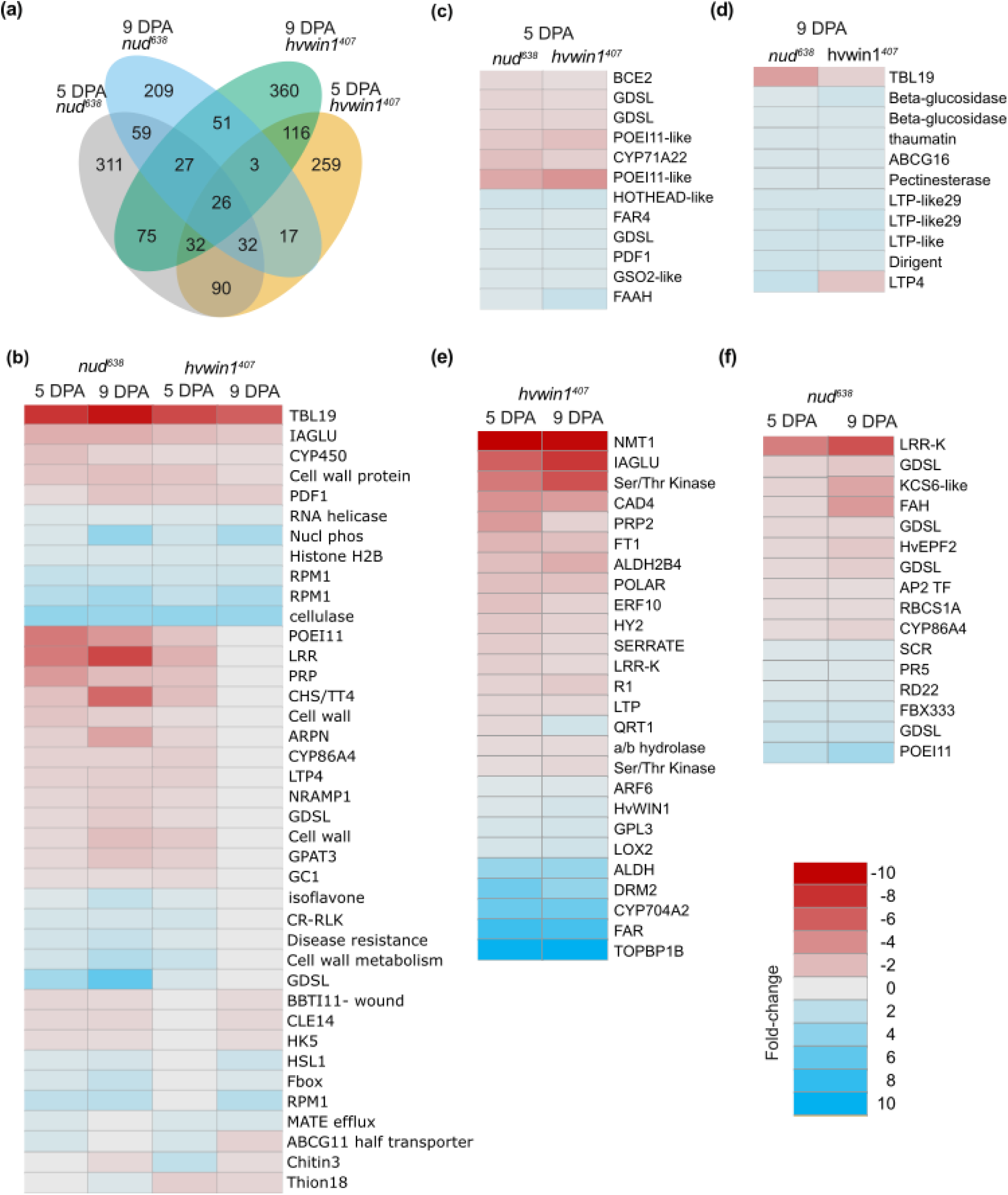
RNA-seq reveals shared and distinct gene expression patterns associated with HvWIN1 and NUD function. (a) Venn diagram describing the overlap between differentially expressed genes (DEGs) between 5 and 9 days post anthesis (DPA) and between in *nud^638^* and *hvwin1^407^* compared to Bowman. (b-e) DEG heat maps showing fold-change in gene expression using a colour scale from upregulated (red) to downregulated (blue). (b) DEGs commonly misregulated at all comparisons (5 and 9 DPAs in *nud^638^* and *hvwin1^407^* compared to Bowman) or in at least three comparisons. (c) DEGs misregulated at 5 DPA in *nud^638^* and *hvwin1^407^* compared to Bowman. (d) DEGs misregulated at 9 DPA in *nud^638^* and *hvwin1^407^* compared to Bowman. (e) DEGs uniquely misregulated in *nud^638^* at both 5 and 9 DPAs. (e) DEGs uniquely misregulated in *hvwin1^407^* at both 5 and 9 DPAs.

## DISCUSSION

The *BDG* gene was initially cloned in Arabidopsis and described as essential for leaf and flower cuticular integrity (Kurdyukov et al., 2006a). Here, we report the first functional characterisation of a *BDG-like* gene outside of dicots. We demonstrate that barley and wheat *BDG* genes control cuticular features, including leaf cuticular integrity and the wax bloom, similar to roles previously described for *HvWIN1* (McAllister et al., 2022), and as shown here for wheat *TdWIN1*. We also report that *HvBDG1* and *HvWIN1* are essential for strong hull to caryopsis adhesion in barley, marking the most significant advance in understanding this trait since the identification of *NUD* almost 20 years ago (Taketa et al., 2008). We show that NUD and HvWIN1 promote *HvBDG1* expression in leaves and grain, which may relate to their control of cuticular features, as well as reveal an essential function for NUD in leaf cuticle integrity. Taken together, our work extends the networks and genetic mechanisms controlling cuticular integrity and specialisations.

We propose that NUD, HvWIN1 and HvBDG1 contribute to a shared hull to caryopsis adhesion pathway. This pathway does not function without NUD, only partially functions without HvBDG1 and/or HvWIN1, and may involve NUD/HvWIN1 induction of *HvBDG1* but not NUD promotion of *HvWIN1* expression. Firstly, *hvwin1^407^, hvbdg1^156^* and *hvbdg1^156^ hvwin1^407^* mutants showed grain skinning rather than naked grain. Secondly, *hvwin1^407^* and *hvbdg1* mutants exhibited intermediate changes to pericarp ultrastructure between wild-type and *nud^638^*, suggesting partial defects in a hull to caryopses adhesion programme. Thirdly, both NUD and HvWIN1 promoted *HvBDG1* expression coincident with hull adhesion, RNA *in situ* hybridisations showed overlapping expression for all three genes in the pericarp epidermis and integuments, and *hvbdg1^156^*’s more severe skinning phenotype was epistatic to *hvwin1^407^*, which collectively supports a downstream role for HvBDG1. Fourthly, the finding that *HvWIN1* expression increases slightly in *nud^638^* caryopses while *NUD* expression remains unchanged in *hvwin1^407^*, suggest NUD likely does not regulate *HvBDG1* and/or hull adhesion by influencing *HvWIN1* levels. Lastly, comparative RNA-seq resolved multiple cuticular and developmental genes sensitive to NUD but not HvWIN1 function and vice versa. Based on these data, we suggest that NUD and HvWIN1 act independently to promote hull to caryopses adhesion, potentially via partially shared targets, such as *HvBDG1*, with NUD playing a more dominant role.

In contrast to the assertion that *NUD* acts exclusively in the grain due to “strict [expression] localized to the testa” (Taketa et al., 2006), we discovered that NUD is essential for leaf cuticle integrity. In fact, leaf permeability increased in *nud^638^ hvwin1^407^* the *hvwin1 nud* double mutantcompared to single mutants, which correlated with a further decrease in *HvBDG1* expression, suggesting HvWIN1 and NUD have an additive relationship in respect to leaf cuticular integrity. Moreover, neither *HvWIN1* nor *NUD* function influences each other’s expression in leaves. Thus, HvWIN1 and NUD may also non-redundantly and independently control leaf cuticle integrity by upregulating *HvBDG1* expression.

HvWIN1 and NUD promotion of *HvBDG1* expression is consistent with SHN factor-driven upregulation of *BDG3* expression in Arabidopsis (Shi et al., 2011). Since *BDG* and *WIN/SHN* genes are co-expressed across land plants (Kong et al., 2020; McAllister et al., 2022; this paper), WIN/SHN regulation of *BDG* expression could be fundamental to cuticle integrity across species. That said, phylogenetic analyses revealed a BDG-like protein in the charophyte *Klebsormidium nitens*, a terrestrial green alga which develops a primitive cuticle, which may indicate that BDG proteins predate land plants (Fig S4). Furthermore, HvBDG1 cannot be solely responsible for *WIN/SHN* control of cuticular integrity since *nud^638^ hvwin1^407^* the cuticles of the *hvwin1* double mutant are more permeable than *hvbdg1^156^* single mutants, suggesting misregulation of other core cuticular metabolism and transport genes must also contribute to compromised cuticular integrity in *nud^638^ hvwin1^407^*. We speculate that *HvBDG4*, moderately expressed in leaf epidermis and stem, could represent such an additional target (Fig S5). Redundancy between *HvBDG*s may also explain skinning rather than naked grain in *hvbdg1^156^* mutants as well as the mild *hvbdg1^156^* phenotypes in barley compared to the severe leaf fusion, deformation and dwarfism of Arabidopsis *atbdg1* mutants (Kurdyukov et al., 2006a; Jakobson et al., 2016). However, SHN genes in barley also have non-overlapping roles, as *nud^638^* mutants do not share HvWIN1’s role in the promotion of cuticular wax blooms. Thus, HvWIN1 likely has an additional function to NUD in these tissues during wax bloom deposition. Taken together, HvWIN1 and NUD play dominant roles in two late stage cuticular elaborations in barley, the wax bloom and hull adhesion, respectively but retain functional, independent overlap in controlling leaf cuticle integrity and hull adhesion, both of which likely involve upregulation of *HvBDG1*.

*HvWIN1*, *NUD* and *HvBDG1* co-expression in the integuments, nucellar epidermis and the pericarp epidermis - three maternal tissues which form cuticles (Freeman and Palmer, 1984) – may indicate that WIN/SHNs promote *BDG* expression in the general context of cuticle formation, even in tissues not exposed to the atmosphere. In support, *AtBDG1* is induced in internal cells that form a *de novo* cuticle as they transdifferentiate into epidermis exposed following floral organ abscission in Arabidopsis (Wen et al., 2025). Post fertilisation, the barley caryopsis grows rapidly, first through longitudinal cell expansion and then widening as the endosperm fills with starch during grain maturation (Brinton et al., 2019). Cuticular material must be deposited in step to maintain nucellar, integument and pericarp layer integrity during early phases of caryopsis growth; however, all maternal tissue layers are substantially modified during grain maturation. For instance, the outer integuments become heavily cuticularized and flatten into the testa following degradation of inner integuments, while the nucellar epidermis crushes into the testa and internal pericarp tissue layers progressively crush against pericarp epidermis during grain expansion (Briggs, 1978). We noted that the pericarp epidermis showed cuticular ridges but at 11 DPA, these ridges sometimes showed thick cuticular plaques or deposits most obvious in Bowman and less so in the *hvbdg1^156^* and *hvwin1^407^* caryopses and not observed in *nud^638^*. Arabidopsis petals also form cuticular ridges that are proposed to help elongating petals slide past with neighbouring organs, as loss or defective ridges lead to petal adhering and folding on itself. Interestingly, both knockdown and overexpression of *SHN* genes in Arabidopsis causes folded petals, with the knockdown associated with reduced petal and sepal ridges, and lowered expression of cutin synthases and *AtBDG3* (Shi et al., 2011), while overexpression caused ectopic ridges, cuticular wax deposits, excess cuticular components and enhanced cuticular permeability (Aharoni et al., 2011). Thus, careful dosage of SHN expression during organ growth and maturation may be important for proper cuticle development. Cuticular ridges in Arabidopsis may reflect a two-step process of early-stage formation and then maintenance of ridges in final length, maturing organs, the latter associated with additional cutin deposition (Li-Beisson et al., 2009; Hong et al., 2017). We speculate that the pericarp cuticle may also develop through a two-step process where early pericarp cuticular ridges may help the caryopsis slide along the encasing hulls and expand along the hull cavity, but at the end of grain elongation, the ridges are modified to promote adhesion to overlying hulls as grains mature and fill with starch. We suggest that the combined activities of NUD and HvWIN1 mediate the switch from non-adherent to adherent via late-stage deposition and/or modification of cuticular material, the ‘cementing layer’ described by Gaines et al. (1986), and the cuticular deposits we observed at 11 DPA, to ensure that the pericarp surface can adhere to the inner hulls. We suggest that NUD and HvWIN1 independent control may suggest that the grass-specific duplication giving rise to separate NUD and HvWIN1 clades (McAllister et al., 2022) may have been co-opted in barley as a mechanism to purposefully generate high and localised doses of SHN activity for sufficient production of cuticular components during late-stage organ maturation leading to compromised and sticky cuticles. We note that HvWIN1 promotion of the wax bloom is also a late stage cuticular modification in maturing organs. Interestingly, Kakeda et al. (2011) reported that transgenic overexpression of *NUD* in rice did not lead to adherent grain but also that *NUD* expression was not maintained in maturing caryopses (Kakeda et al., 2011), suggesting that barley may have a species-specific mechanism to maintain *NUD* expression in late-stage caryopses. It would be interesting to examine the regulatory regions of these two clades across grasses for signatures of barley-specific regulatory motifs.

Studies on cuticular ridges also suggest that cuticle chemistry interacts with cuticular ultrastructure to influence adherent properties. Ridges are associated with cuticle buckling driven by cuticular material properties characterised by a relatively stiff outer ‘cuticle proper’ and a softer inner cuticular layer, which often stains darkly under TEM, that may be continuous with the cell wall (Martens, 1933; Li-Besson et al., 2019; Kourounioti et al., 2012; Kourounioti et al., 2013; Huang et al., 2017; Mazurek et al., 2017; Airoldi et al., 2021). In addition, multiple organ fusion mutants also demonstrate excess cuticular component accumulation, inferred to reflect perceived cuticle damage (Lolle et al., 1998; Voisin et al., 2009; Mazurek *et al*., 2017; Huang et al., 2023). We show that at 7 DPA the thick Bowman pericarp cuticles had darkly stained electron-lucent globules and fibrillar extensions at the cell wall-cuticle interface (Fig S12) which itself became progressively less distinct by 11 DPA (Fig 4e), possibly indicating cuticular layer merging with the underlying cell wall. We are intrigued by the similarities between the Bowman cuticular ultrastructure and the Arabidopsis *cyp77a6* mutant petal cuticles described to have a thick amorphous cuticle with little distinction between the cell wall and fibrous cell wall-like material, defective cuticular integrity and extremely adherent petal surfaces (Mazurek et al., 2017). Brennan et al. (2019) observed similar interfaces in several barley cultivars undergoing hull adhesion which they speculated could reflect pectic polysaccharides, consistent with cell wall changes. In contrast, *nud^638^* cuticles were thinner, lacking a cuticle proper/cuticular layer distinction, with a very sharp boundary to the cell wall without globules or fibrillar extensions, while intermediate phenotypes in skinning mutants, including globule accumulation restricted to the cell wall in *hvbdg1^156^*, suggesting an alteration in the normal changes observed in covered barley pericarp cuticle and cell wall interface. Arabidopsis *bdg* mutants develop wall deformations including fibrous networks between wall and cuticle (Voisin et al., 2009), also indicative of BDG roles in this interface. Altogether, the changes in the cuticle cell-wall interface may be integral to the adherent properties of the late-stage pericarp cuticle.

Consistent with high SHN expression leading to cuticle permeability, our previous work demonstrated that at 11 DPA covered barley’s pericarp epidermal layer tears upon hull removal and is easily penetrated by dyes (Campoli et al., 2024). Intriguingly, *nud^638^* pericarps remain intact in mature grain with permeability to dyes or surface lesions (Fig S18), and show increased levels of sitosterol, a component associated with enhanced cuticle integrity (Chen et al., 2020a). We found an association between strong hull adhesion and surface chemistry shifts from short to longer chain fatty acids, as well as no increase in C18 fatty acids in *nud^638^*, in contrast to the other lines, pointing to a possible role in adherent properties, broadly supporting previous work that skinning cultivars have fewer cuticular fatty acids compared to cultivars with better adhesion (Brennan et al., 2019). Interestingly, increased C16 and C18 fatty acids in floral organ mutants in Medicago corresponded to adhesion in leaves and flowers (Wang et al., 2023). We suggest that these changes could indicate a role for these lipid components in the loss of adhesion in naked and skinning grain. Despite obvious cuticular thickening in the outer pericarp, we did not detect increased cutin components coincident with hull adhesion nor different cutin amounts in the hull adhesion mutants with thinner pericarp cuticles. However, our methods do not detect possible differences in cutin polymerisation, which Arabidopsis BDGs may help catalyse (Kurdyukov et al., 2006a). Another option could be that our extraction methods did not capture cutin or other components in other tissues which may contribute to adherent properties. For example, transfer of lipid components onto the cell surface from another layer occurs during pollen coat formation wherein a mixture of tapetum and endothecium-derived lipids, including VLFCAs and cutin are deposited onto the pollen wall surface making it sticky, a process driven by early and late regulatory cascades in anthers involving specific MYBs and downstream KCSs (Bai *et al*., 2019; Battat *et al*., 2019; Chen *et al*., 2020b; Zhang *et al*., 2021a,b). Strong integument expression of *NUD, HvWIN1* and *HvBDG*, as well as the progressive crushing of the pericarp, make us wonder if the integuments may produce the cementing layer components ultimately found on the weakened and permeable pericarp epidermis.

Despite advancing our understanding of HvBDG1’s role in cuticle development in barley, its exact molecular function in this process remains poorly understood. HvBDG1 could function directly in synthesising lipid precursors for cuticle development or supporting fatty acyl chain elongation, with a reduction in specific substrates in *hvbdg1^156^* inhibiting normal trafficking processes. Given that *HvBDG1* encodes a hydrolase, it may be involved in cleavage of fatty acyl precursors for subsequent reactions. Initial characterisation of barley *cer-a* alleles and their role in β-diketone synthesis showed a similar trend to our results, with reductions in β-diketones and other ubiquitous waxes (von Wettstein-Knowles, 1976). von Wettstein-Knowles proposed a role for *cer-a* upstream of these processes, before their divergence (von Wettstein-Knowles, 2017). In this case, we would expect HvBDG1 to function intracellularly rather than extracellularly. Kurdyukov et al. (2006a) could not definitively confirm the localisation of AtBDG1 due to the limited resolution of their method, therefore an intracellular role for HvBDG1 is plausible and merits further investigation. We speculate that the grass-specific N-terminal region could also be relevant to these differences in localisation pattern.

In conclusion, our work greatly expands the genetic networks and molecular activities important for cuticle development in barley and our understanding of shared and species-specific cuticular specialisations in plants.

## ACKNOWLEDGEMENTS

We are deeply grateful to the NordGen seedbank, National Plant Germplasm System at the U.S. Department of Agriculture and Okayama University for barley germplasm. We are also deeply grateful to the Barley Genetics group at the James Hutton Institute for providing germplasm and expertise, especially Pauline Smith, Richard Keith, Chris Warden, and Dr Joanne Russell. We thank the John Innes Centre Germplasm Resources Unit, a National Bioscience Research Infrastructure supported by the UKRI-BBSRC, grant number BBS/E/JI/23NB0001 for conserving and supplying germplasm through www.seedstor.ac.uk and the Niab Glasshouse Team for wheat plant growth support, facilities access and expertise. We also acknowledge services and support from the Genome Technology Lab at the JHI as well as guidance from Dr Petra Boevink in the localisation experiments and Dr Alexandre Foito in generation of GC-MS data. We are also grateful to Ms. Claire Traynor and the Mylnefield Lipid Analysis team (James Hutton Ltd., Dundee, Scotland) for access to facilities and expertise in GC-FID and GC-MS. We acknowledge the guidance and technical support of Dr Raymond Campbell and Dr Kyriakos Varypatakis for running the *in situ* hybridisation experiment. Finally, we are grateful for support from the University of Dundee Imaging Facility as well as Imaging Technologies at the JHI.

