## Supplemental Notes for "Barley BODYGUARD controls cuticular specialisations regulated by SHINE transcription factors"

**Supplemental Note 1**

We generated protein models for wild-type Bonus HvBDG1 using ReFOLD-refined IntFOLD-TS structural prediction with AlphaFold2 and trRosetta2. We observed that ReFOLD refinement improved the prediction by 2.1% resulting in a DeepUMQA-X model quality assessment with TM score of 0.865 and Global- Inter-residue distance deviation (lDDT) of 0.718 (Fig 1). Global lDDT is brought down by an ≈ 76-128aa Nʹ terminal region consisting of unstructured glycine-rich linkers, two, short alanine-rich α-helices and a serine-rich region that may provide scaffolding for molecular recognition and adopt a stable structure upon binding a target protein/substrate or environmental changes.

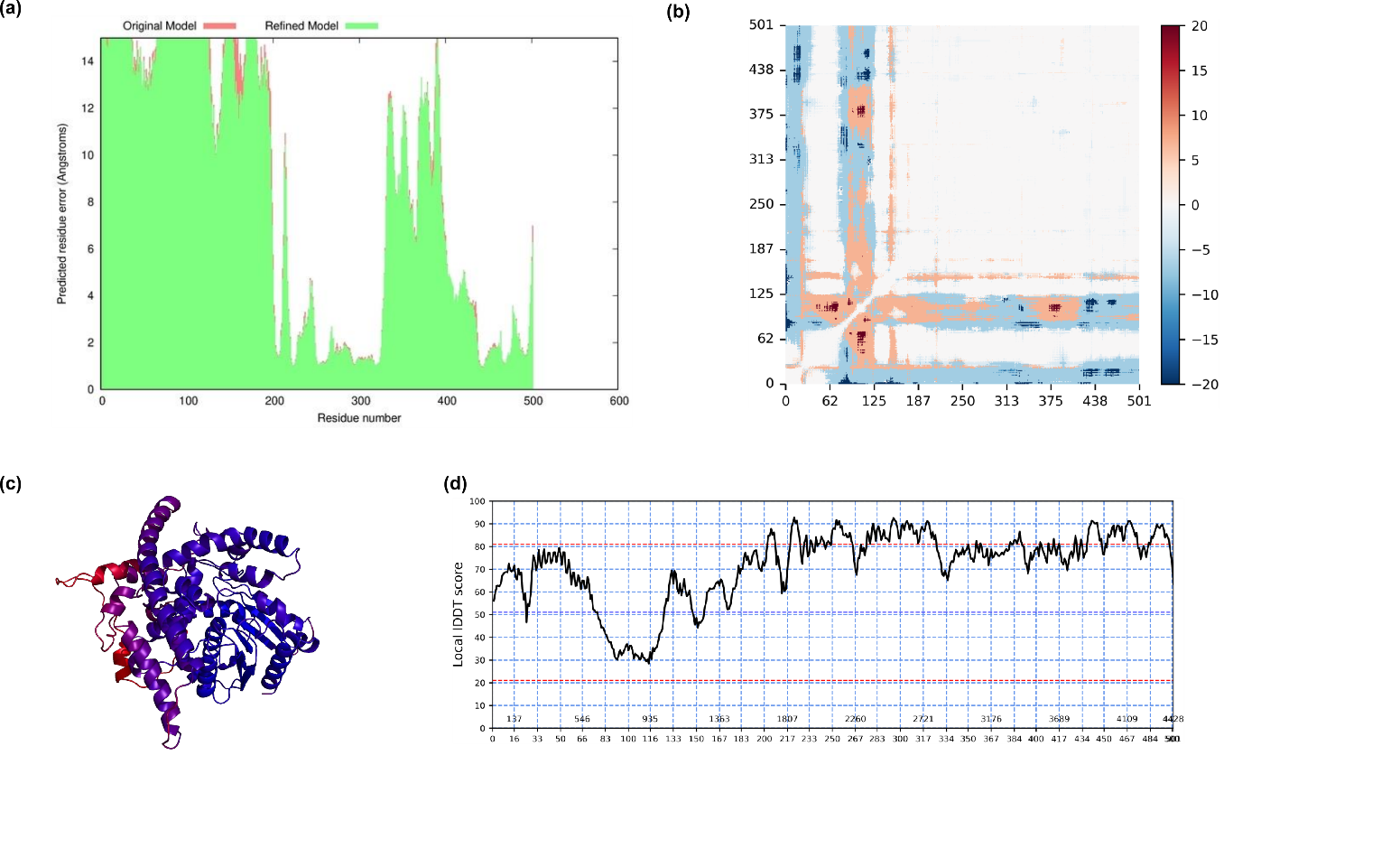

**Figure 1. Model quality assessments.** (a) Predicted residue error (Å), for the original (orange) and refined (green) models. (b) Inter-residue distance deviation (IDDT) from this assessment. (c) the lDDT mapped onto the model prediction using a low-high/red-blue scale and D shows the per-residue atomic lDDT.

To assess stereochemical quality, we conducted Ramachandran plot analysis based on a reference set of 118 high-resolution structures (resolution ≤ 2.0 Å, R-factor ≤ 20%) (Fig. 2). The analysis revealed that 91.9% of residues (406 amino acids) were located within the most favoured regions (A, B, L), while 7.9% (35 residues) occupied additionally allowed regions (a, b, l, p). No residues were found in generously allowed regions (~a, ~b, ~l, ~p), and only one residue (0.2%) was located in a disallowed region. This outlier corresponds to S299, a catalytically important residue, suggesting that its strained backbone conformation may facilitate stabilization of the reaction intermediate. Additional residues with unfavourable Ramachandran Z-scores (< 3.00) include I18, L265, S122, V116, V284, and V463, potentially reflecting local conformational flexibility or functional constraints.

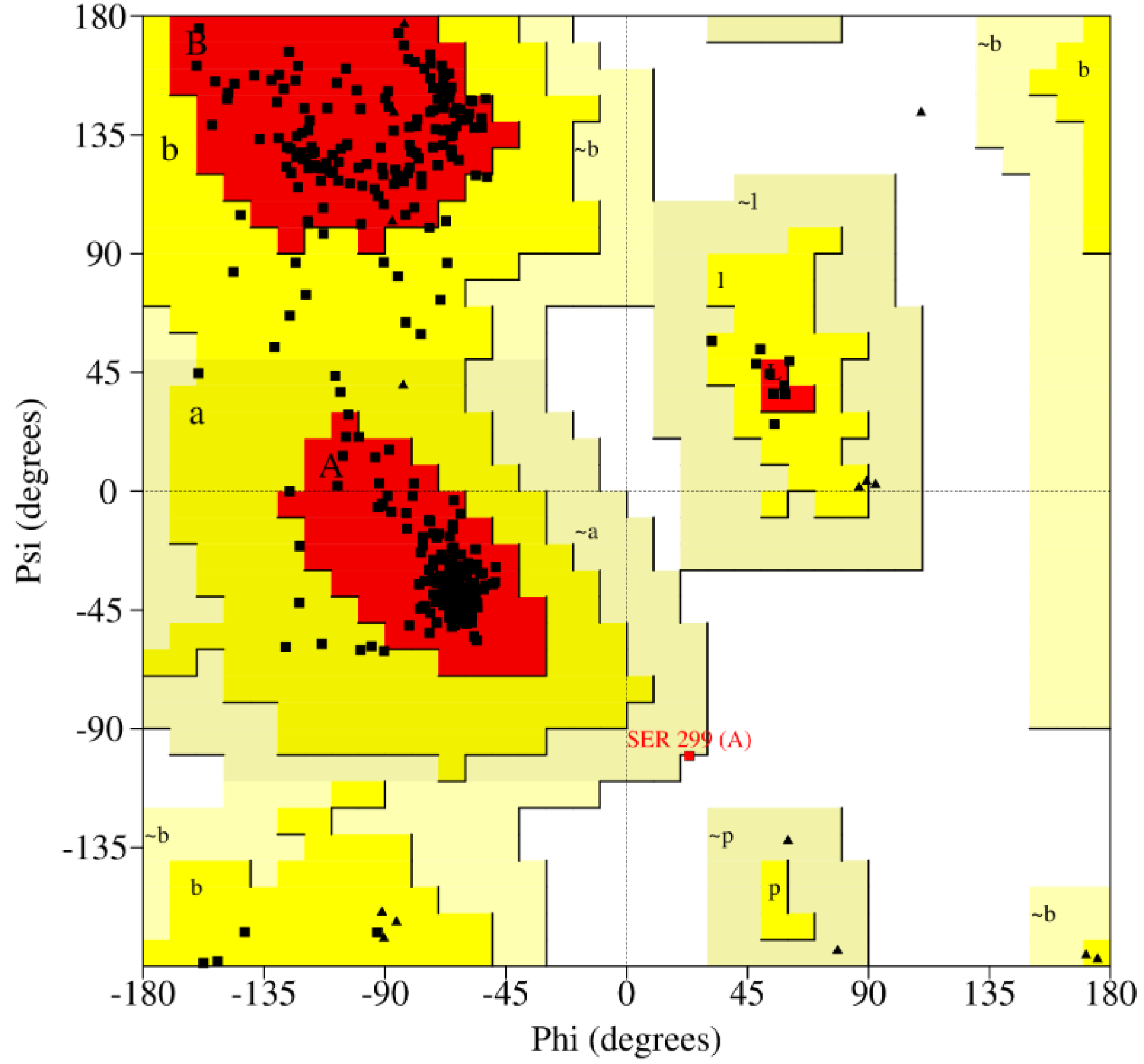

**Figure 2: Ramachandran Plot Analysis of Bonus HvBDG1.** Regions are coloured based on most favoured (red), additional allowed (yellow), generously allowed (light yellow) and disallowed (white). The catalytically active residue S299 is highlighted in red.

We next generated protein models for the mutant HvBDG1^156^ H407/R protein, again using ReFOLD-refined IntFOLD-TS structural prediction utilising AlphaFold2 and trRosetta2. In this case, ReFOLD refinement improved the prediction by 2.0% resulting in a DeepUMQA-X model quality assessment with TM-score of 0.840 and Global-lDDT of 0.667 (Fig 3). Global lDDT was brought down in the same manner as the wild-type Bonus prediction by an ≈ 76- 128aa Nʹ terminal region.

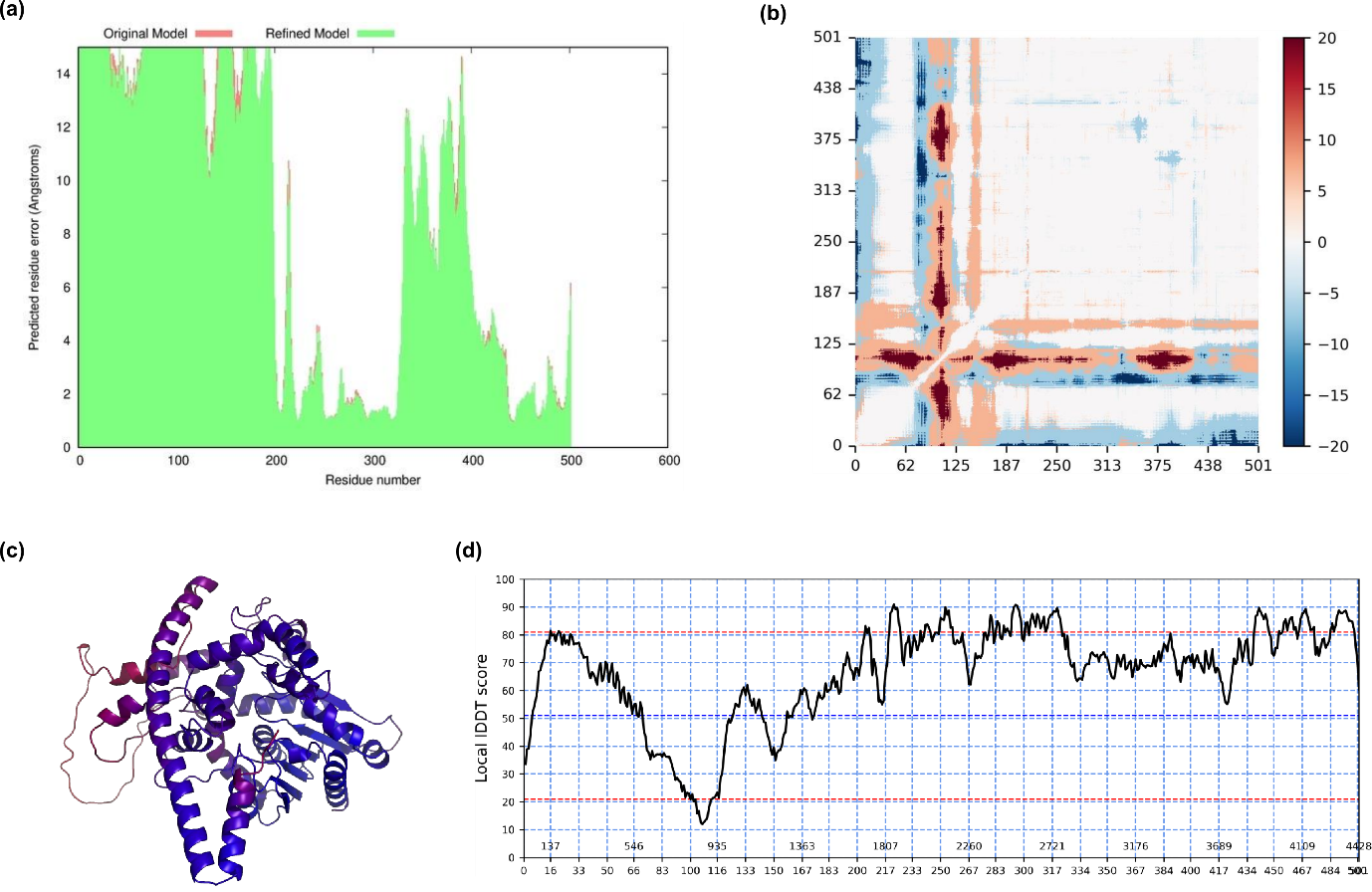

**Figure 3. Model quality assessments of HvBDG1^156^ H407/R.** (a) Predicted residue error (Å), for the original (orange) and refined (green) models. (b) Inter-residue distance deviation (IDDT) from this assessment. (c) the lDDT mapped onto the model prediction using a low-high / red-blue scale and D shows the per-residue atomic lDDT.

**Supplemental Note 2**

*Identification of durum wheat BDG1*

Using barley *BDG1* coding regions (CDS) from gene model transcript *HORVU.MOREX.r3.7HG0644300.1* as a query for BLASTn analysis of the durum wheat *cv*. Svevo reference genome (assembly Svevo.v1; Maccaferri et al. 2018) using Ensembl Plants (Harrison et al., 2024), identified two genomic regions with e-values <e-^27^, corresponding to the positions of two gene models: *TRITD7Av1G017740* (termed here *TdBDG1-A1*) on the short arm of chromosome 7A at 32.1 Mb (longest High-scoring Sequence Pair: length = 331, 2.5e-^161^) and *TRITD4Av1G244120* (*TdBDG1-A2*) on the long arm of chromosome 4A at 685.5 Mb (longest High-scoring Sequence Pair: length = 331, 2.5e^-161^). Equivalent BLASTn search of the bread wheat *cv*. Chinese Spring genome (assembly RefSeq v1.0; IWGSC, 2018) identified three genomic regions with e-values <e-^27^, corresponding to the positions of three gene models: *TraesCS7A02G068800* on the short arm of chromosome 7A at 34.9 Mb (longest High-scoring Sequence Pair: length = 331, 3.4e^-161^), *TraesCS4A02G419700* on the long arm of chromosome 4A at 690.5 Mb (longest High-scoring Sequence Pair: length = 331, 3.4e^-161^), and *TraesCS7D02G063500* on the short arm of chromosome 7D at 34.5 Mb (longest High-scoring Sequence Pair: length = 331, 5.7e^-166^). The translocated bread wheat 7B chromosomal segment known to be homoeologous to bread wheat chromosomal regions spanning 0-75 Mb on 7A and 0-60 Mb on 7D (Zhou et al., 2020) span the locations of *TaBDG1-A* (7A at 35.9 Mb) and *TaBDG1-D* (7D at 34.351 Mb) are found, respectively, confirming the reciprocal chromosomal translocation T(4AL;7BS)1 present hexaploid wheat (Zhou et al., 2020) and durum wheat (Dvorak et al., 2018) resulted in relocation of the ancestral 7B homoeologue to the long arm of chromosome 4A.
