## Supplementary Figures for "Barley BODYGUARD controls cuticular specialisations regulated by SHINE transcription factors"

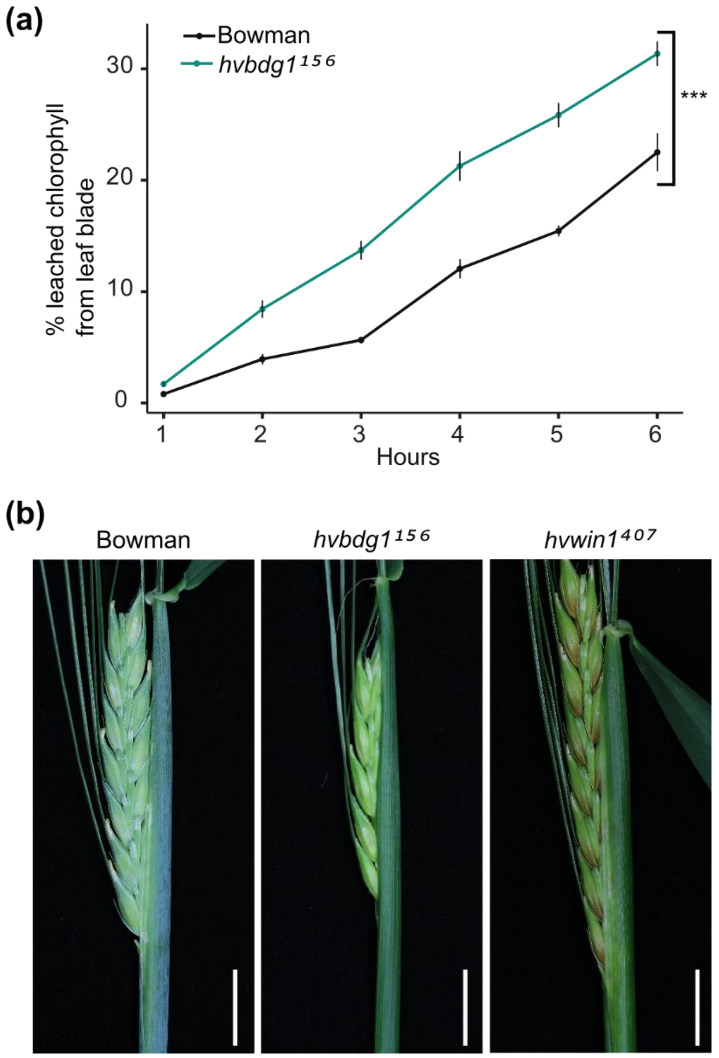


**Figure S1 – Additional BW-NIL phenotypes**. (a) Chlorophyll leaching assay on detached second leaf blades from Bowman and *hvbdg1*$^{156}$ (n = 4, ***$p < 0.001$). (b) Wax bloom phenotypes on leaf sheaths and spikes of in Bowman, *hvbdg1*$^{156}$ and *hvwin1*$^{407}$.

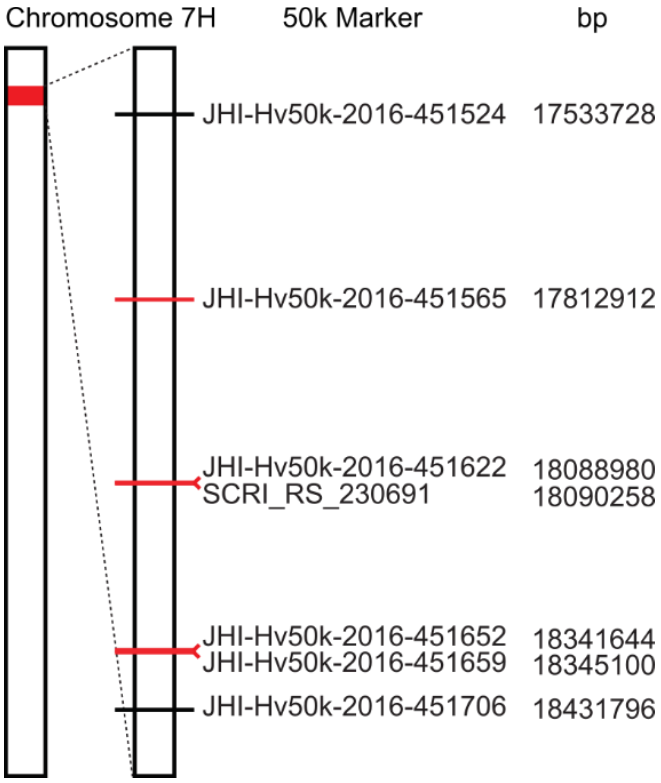


**Figure S2 – Mapping the introgression locus of BW156/*hvbdg1*$^{156}$**. Schematic representation of chromosome 7H of BW156 highlighting the introgression from Bonus in red. In the blow-up, polymorphic markers (red for Bonus, black for Bowman) from the 50k iSelect SNP chip are shown with their positions on the chromosome.

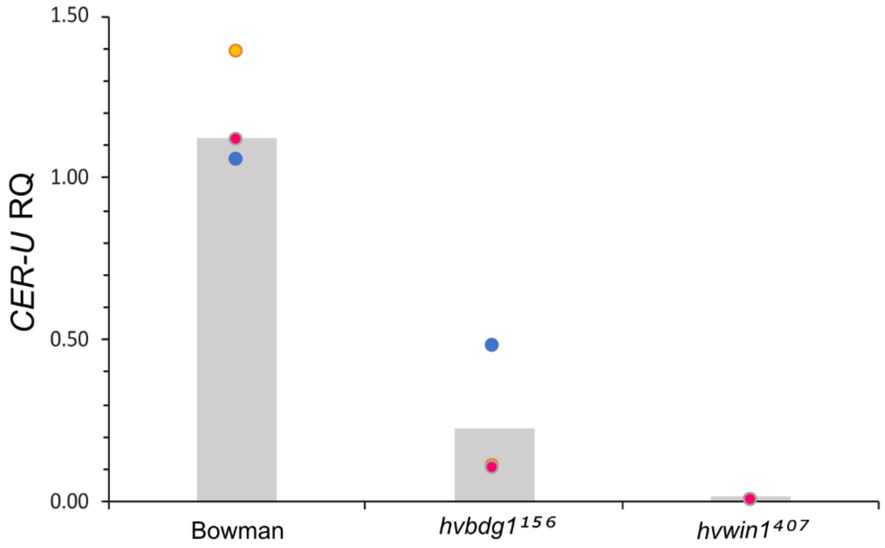


**Figure S3 – *Cer-U* expression in BWNILs mid-flag leaf sheath.** qRT-PCR of *CER-U* transcript levels in Bowman, *hvbdg1*$^{156}$ and *hvwin1*$^{407}$ expressed as relative quantity (RQ). Bars indicate the mean of three biological replicates. Coloured circles show the average of three technical replicates of each independent biological replicate. Flag leaf sheaths were harvested when auricles were 1 to 3 cm above the auricles of the second-to-flag leaf sheaths. (n = 3).

(a)

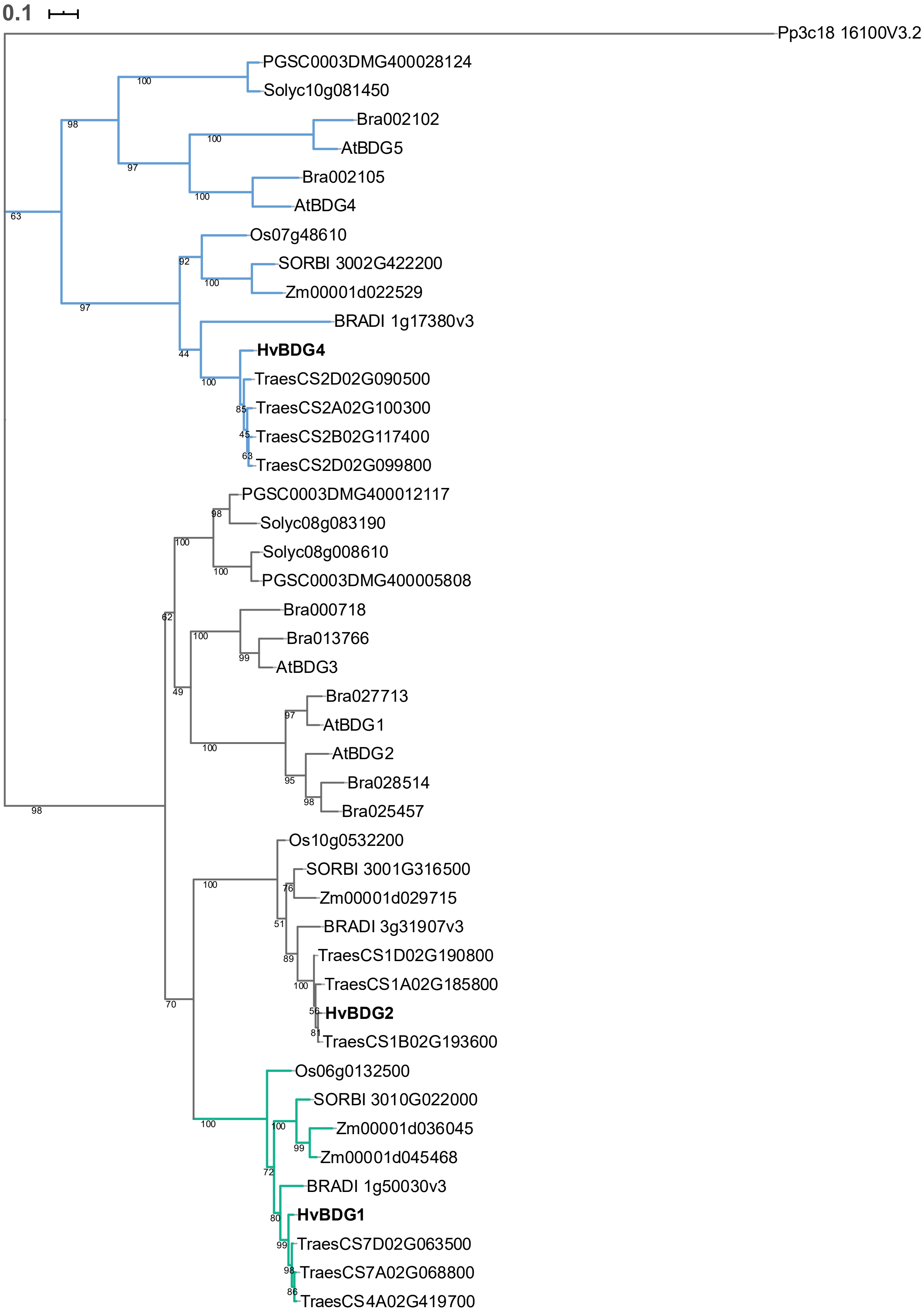
0.1
Pp3c18 16100V3.2
PGSC0003DMG400028124
Solyc10g081450
Bra002102
AtBDG5
Bra002105
AtBDG4
Os07g48610
SORBI 3002G422200
Zm00001d022529
BRADI 1g17380v3
HvBDG4
TraesCS2D02G090500
TraesCS2A02G100300
TraesCS2B02G117400
TraesCS2D02G099800
PGSC0003DMG400012117
Solyc08g083190
Solyc08g008610
PGSC0003DMG400005808
Bra000718
Bra013766
AtBDG3
Bra027713
AtBDG1
AtBDG2
Bra028514
Bra025457
Os10g0532200
SORBI 3001G316500
Zm00001d029715
BRADI 3g31907v3
TraesCS1D02G190800
TraesCS1A02G185800
HvBDG2
TraesCS1B02G193600
Os06g0132500
SORBI 3010G022000
Zm00001d036045
Zm00001d045468
BRADI 1g50030v3
HvBDG1
TraesCS7D02G063500
TraesCS7A02G068800
TraesCS4A02G419700

**(b)**

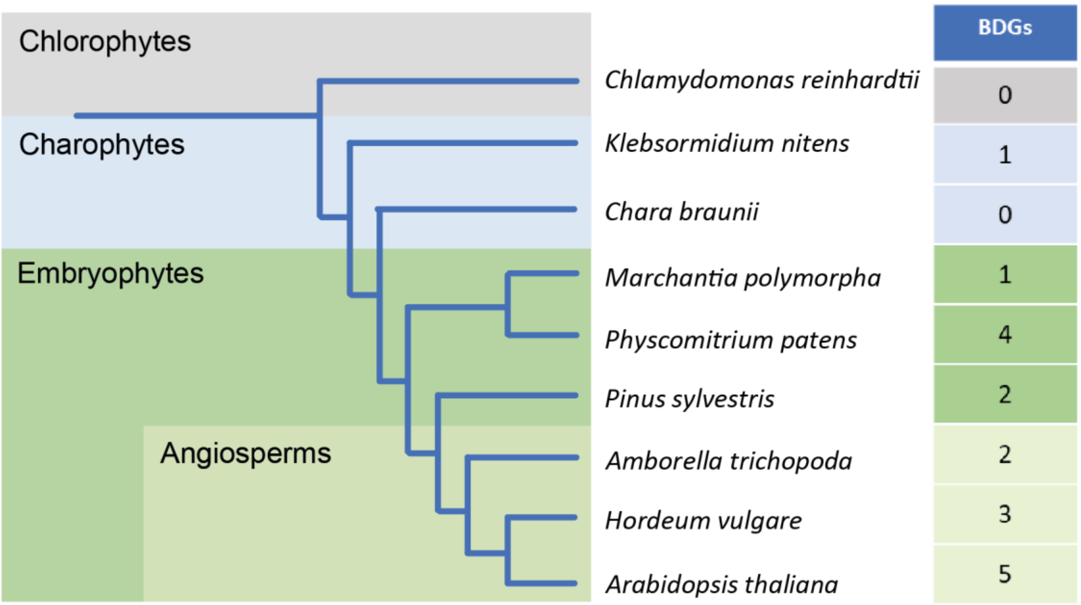


**Figure S4 - Phylogenetic relationship of the BDG family.** (a) Between members of the BDG family in model angiosperm species. Evolutionary analysis was inferred by using the Maximum Likelihood method and JTT matrix-based model in MEGA X (Jones et al., 1992; Kumar et al., 2018). The tree with the highest log likelihood (-12129.36) is shown. The percentage of trees in which the associated taxa clustered together in 500 bootstrap replications is shown next to the branches. A BDG-like protein from *Physcomitrium patens* was used as an outgroup to root the tree. Green: Cereal specific clade of BDG1 proteins. Blue: BDG4 clade diverged before the emergence of angiosperms. Bold: barley proteins. (b) BDG proteins emerged early in the green plant lineage. Schematic of evolutionary relationships between representative species in the green plant lineage and the number of BDG proteins identified in each species.

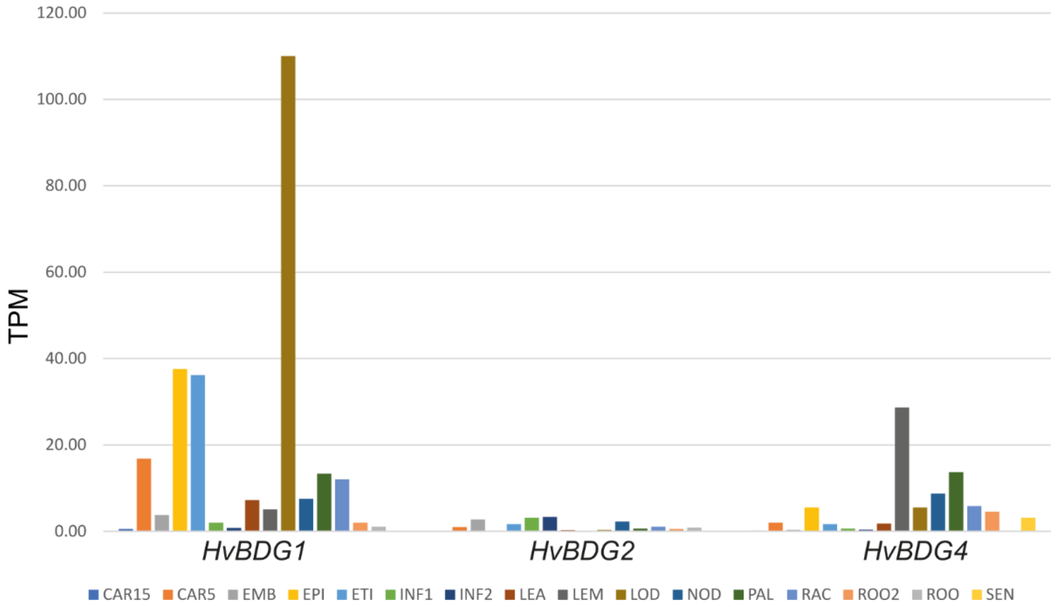


**Figure S5 - Expression profiles of *HvBDG* genes in barley cultivar Morex.** Data extracted from the Barley Expression Database EoRNA (Milne et al., 2021). CAR: caryopsis, EMB: embryo, EPI: epidermis, ETI: etiolated seedling, INF: inflorescence, LEA: leaf, LEM: lemma, LOD: lodicule, NOD: stem, PAL: palea, RAC: rachis, ROO: root, SEN: senescing leaf. Bars represent the mean of three biological replicates. TPM: Transcripts per million.

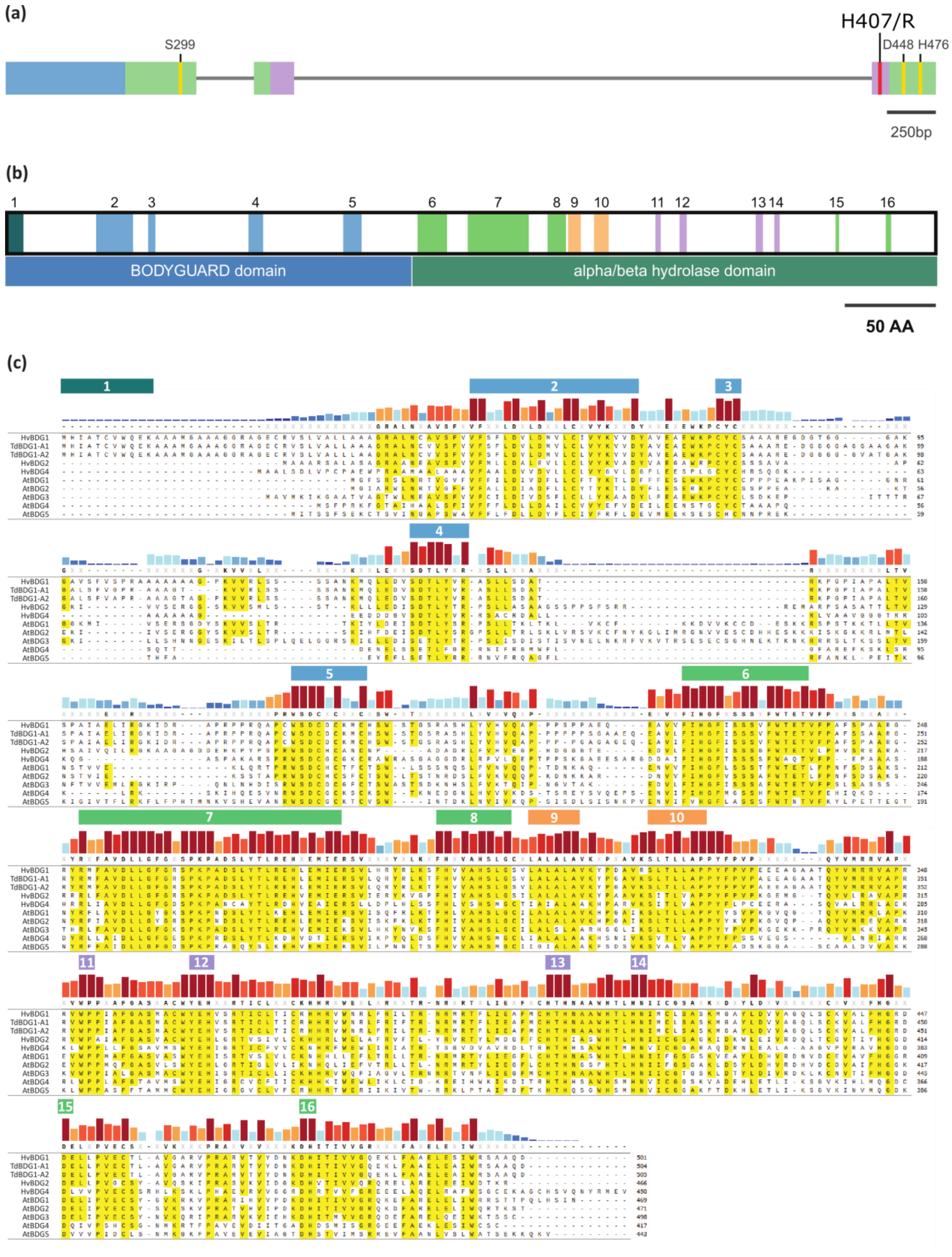


**Figure S6 – BDG protein motifs** (a) Domains mapped onto the *HvBDG1* gene model. Blue: BDG domain. Green: α/β hydrolase core domain. Red: Histidine 407 (H) to Arginine (R) SNP found in BW156 and BW406. Purple: "Lid" domain. Yellow: Amino acid residues of the catalytic triad (Serine 299, Aspartic acid 448 and Histidine 476). (b) BDG motifs plotted onto the HvBDG1 protein. (c) Alignment of characterised BDG proteins with conserved motifs highlighted above. First 3 sequences in alignment belong to the grass specific BDG1 clade containing the conserved N-terminal sequence (red, motif 1). Blue: BDG domain motifs. Green: known alpha/beta hydrolase domain motifs. Orange: novel alpha/beta hydrolase motifs. Purple: novel motifs in the lid domain.

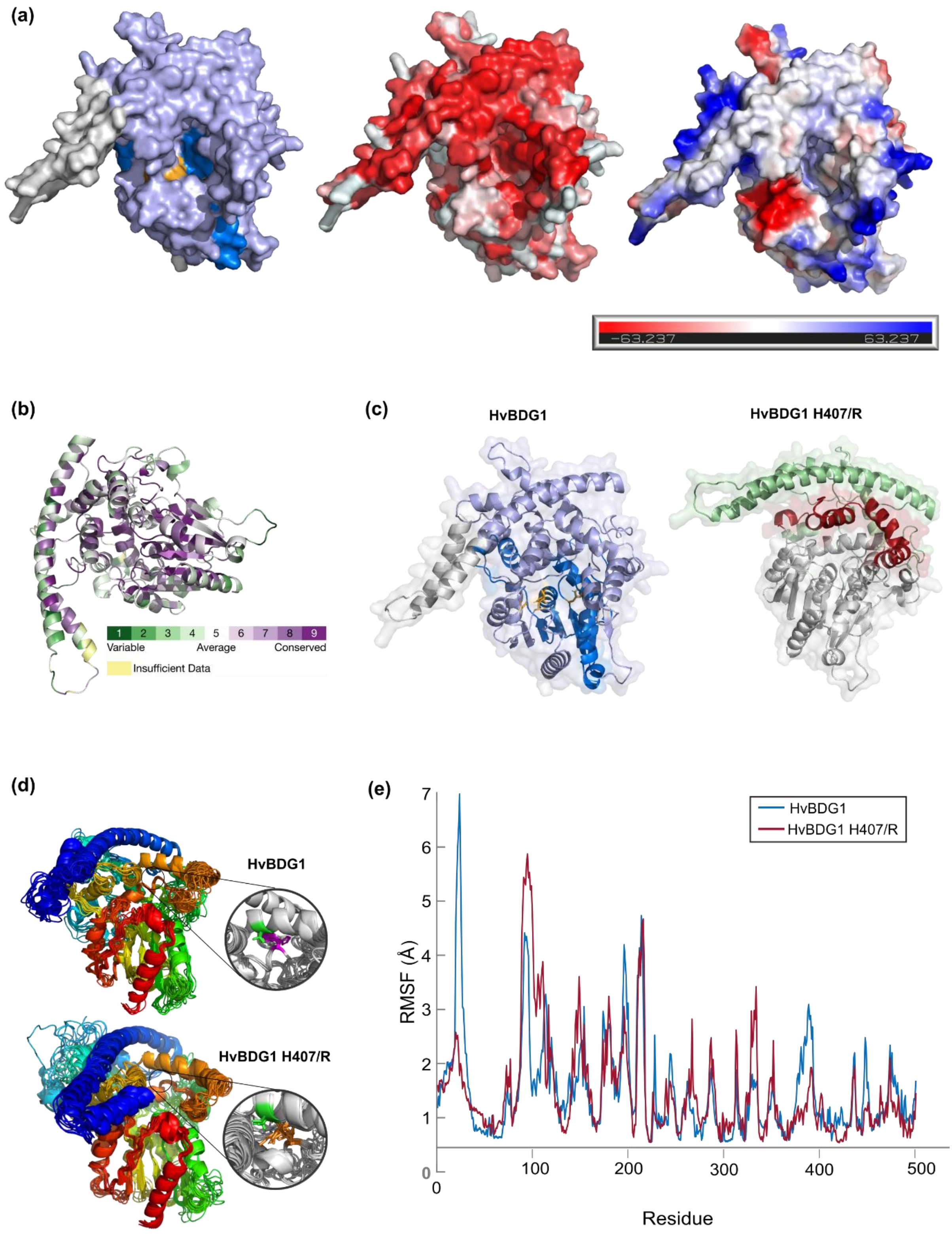


**Figure S7 – Protein modelling of HvBDG1.** (a) HvBDG1 cv Bonus protein model where regions and domains are overlaid with an opaque surface. Model on left coloured by domain: The alpha-beta hydrolase_1 domain (Cdvist ID by hmmer3 and database 30.0, Score: 75.5, region: aa219-347) shown in dark blue within a larger BDG1 domain containing hydrolase in pale blue (NCBI Conserved Domain Database ID, PLN03087, accession: cl30400, region: 42-496, E-value: 0e+00). The catalytic residues S299, D448 and H476 are shown in orange as is the H225, situated in the active site and likely important for stabilising the reaction. Middle panel shows the surface according to Eisenberg's scale of hydrophobicity showing a hydrophobic pocket (red) in which the catalytic residues reside. Right panel presents a qualitative representation of electrostatic potential generated with vacuum electrostatics showing a shallow negatively charged cleft leading to the active site, with a moderately positive charge in the active site pocket. (b) ConSurf Colour-Coded conservation based on MSA of 150 sequence selected using HMMER and UniRef90 with a cutoff of E = 0.0001, a CD-Hit maximum cutoff of 95% and minimum 35% with 10% maximal overlap between homologues, projected on the structural model of Bonus HvBDG1. (c) Ribbon diagrams of HvBDG1 cv Bonus and HvBDG1 H470/R protein models. HvBDG1 cv Bonus model on left shows regions and domains identified by multiple tools. Conserved domain analysis revealed a central Abhydrolase_1 domain (CDvist ID; HMMER3, database v30.0; score: 75.5) spanning residues 219-347, depicted in dark blue. This domain lies within a broader Bodyguard 1 hydrolase domain (NCBI Conserved Domain Database ID: PLN03087, accession: cl30400; residues 42-496; E-value: 0.0), shown in pale blue. Substantial overlap was observed with additional domain annotations, including the UniProt-assigned AB hydrolase-1 domain (residues 221-407), the PANTHER-assigned hydrolase domain (PTHR43689; residues 141-454), and InterPro entries for alpha/beta hydrolases: the superfamily SSF53474 (IPR029058; residues 162-450) and the CATH-Gene3D integrated region (residues 160-461). Overlapping regions are not highlighted. Three putative catalytic residues S299, D448, H476, and a fourth stabilising residue H225 are conserved and indicated in orange. The image on the right shows the approximate region destabilised by site 407H mutation (red), corresponding with the "lid" domain, and the region attached that is predicted to structurally rely upon the position of this (green), corresponding with most of the BDG domain. However, this region includes α-helix 2. We are not confident that this helix interacts with the region in red as we predicted a transmembrane domain within this structure using Phyre2. (d) Protein structural flexibility by coarse-grained protein modelling with CABS-flex 2.0 using ReFOLD-refined Intfold7 structural predictions. Top panel shows the flexibility predictions of the HvBDG1 cv Bonus where histidine aa407 beyond the N' terminus of α-helix 21 is held in place by an interaction internal to α-helix 18 with serine aa368. Bottom panel the same region in the BDG1 H407/R mutant, where the arginine residue no longer interacts with S368 leading to no interaction between α-helix 18 and 21 and a change in conformational state in the local areas area. (e) By residue structural flexibility in HvBDG1 cv Bonus and HvBDG1 H407R. Increased flexibility in the preceding ≈ 150aa and decrease in flexibility in the immediate area between ≈ 375-400aa due to apparent compression of the local environment.

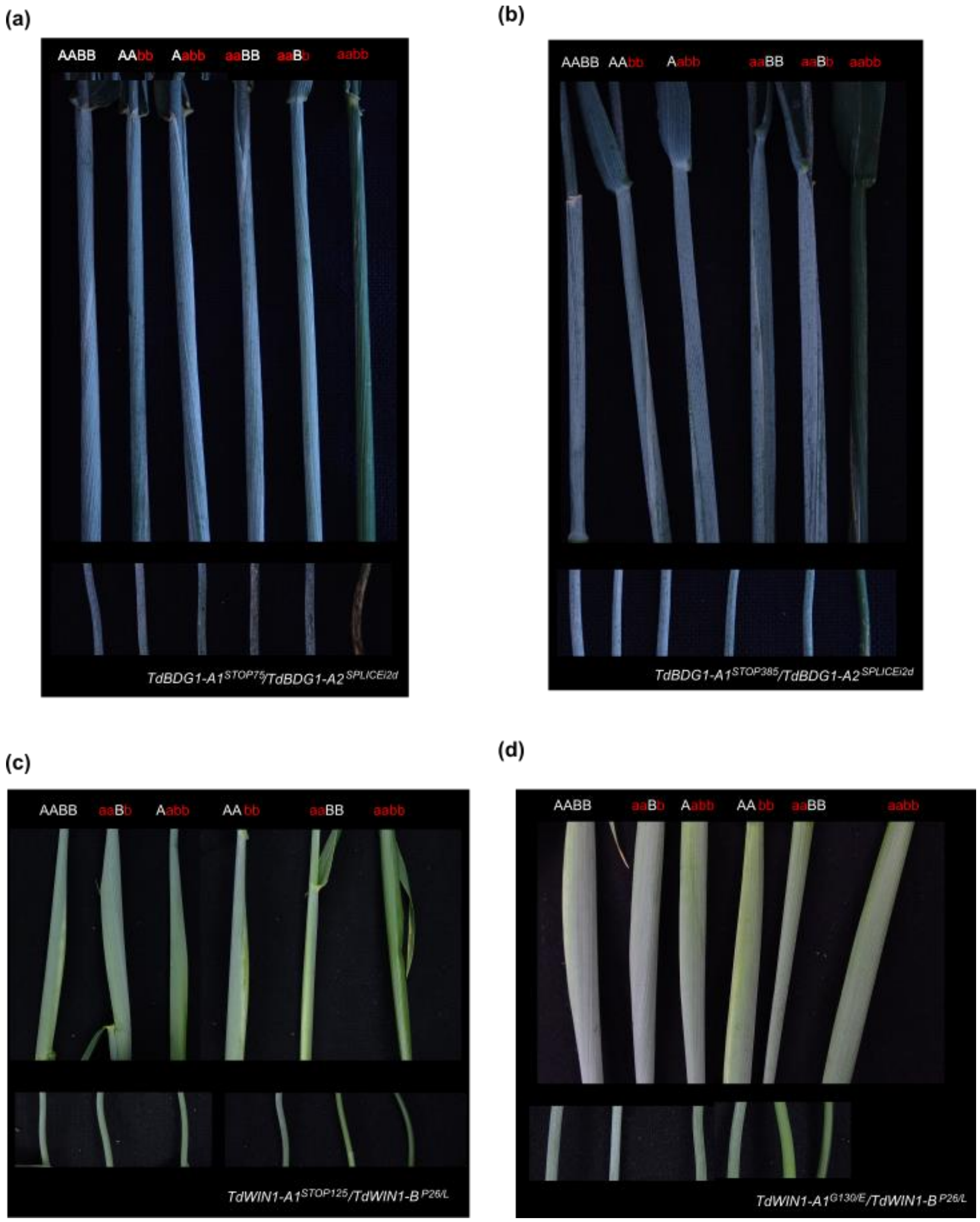


**Figure S8 Wheat stem and sheath phenotypes in *TdBDG1* and *TdWIN1* mutants.** Examples of wax bloom phenotype observed in durum wheat leaf sheath (top row) and stems (bottom row) from $F_2$ individuals carrying different wild-type and mutant alleles at homoeologues of (a,b) *TdBDG1* and (c,d) *TdWIN1*. Two combinations of mutants are shown for both *TdBDG1* and *TdWIN1*. For each combination, allelic state at the A- and B- sub-genome homoeologue is indicated via uppercase (wild-type) or lowercase (mutant) letters, such that homozygous wild-type (AA or BB), homozygous mutant (aa or bb), and combinations of homozygous and wild type (e.g., AABb) alleles are indicated.

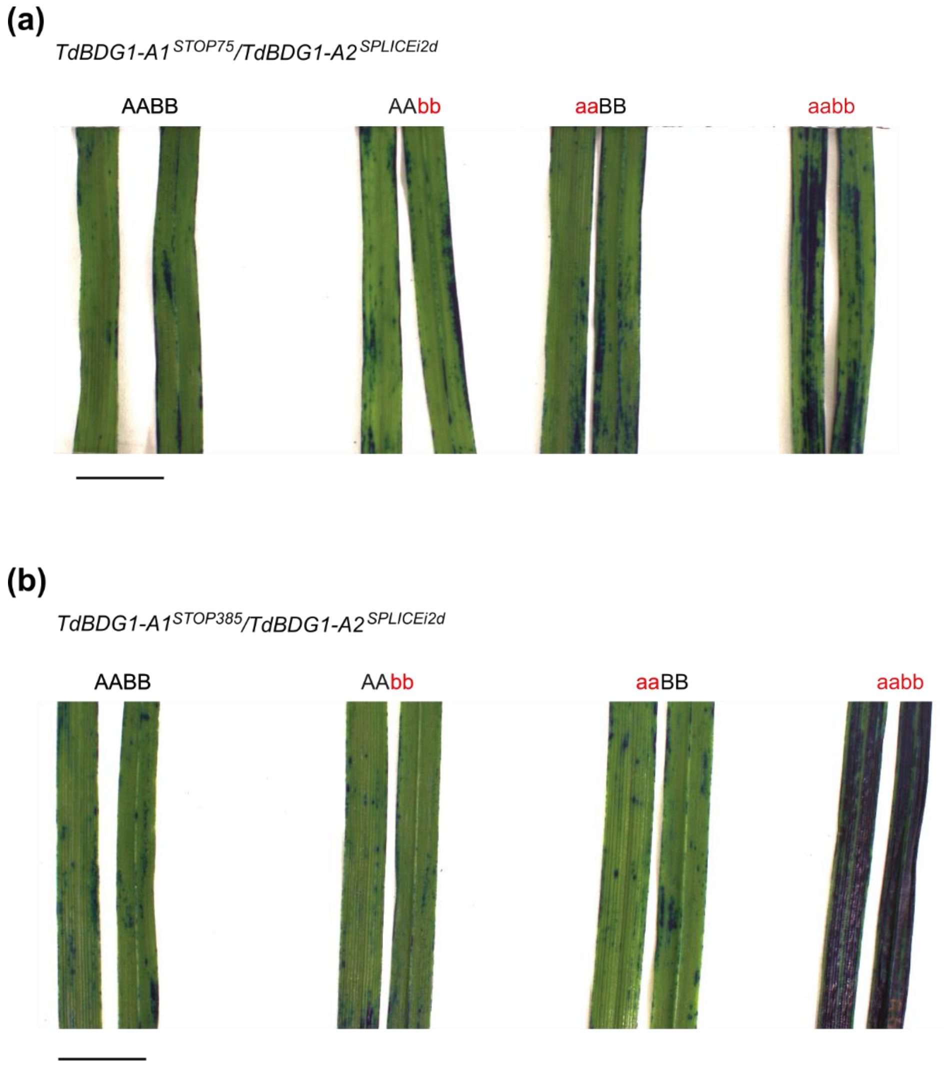


**Fig S9. Wheat leaf blades cuticle permeability.** Detached second leaf blades from wild-type and mutant alleles at homoeologues of (a) *TdBDG1* and (b) *TdWIN1* were submerged in 0.05% Toluidine Blue (TB) for 5 hours. For each genotype, allelic state at the A- and B- sub-genome homoeologue is indicated via uppercase (wild-type) or lowercase (mutant) letters, such that homozygous wild-type (AA or BB), homozygous mutant (aa or bb), and combinations of homozygous and wild type (e.g., AABb) alleles are indicated. Five bioreplicates of each genotype were stained, with two representatives photographed to show adaxial and abaxial sides, respectively.

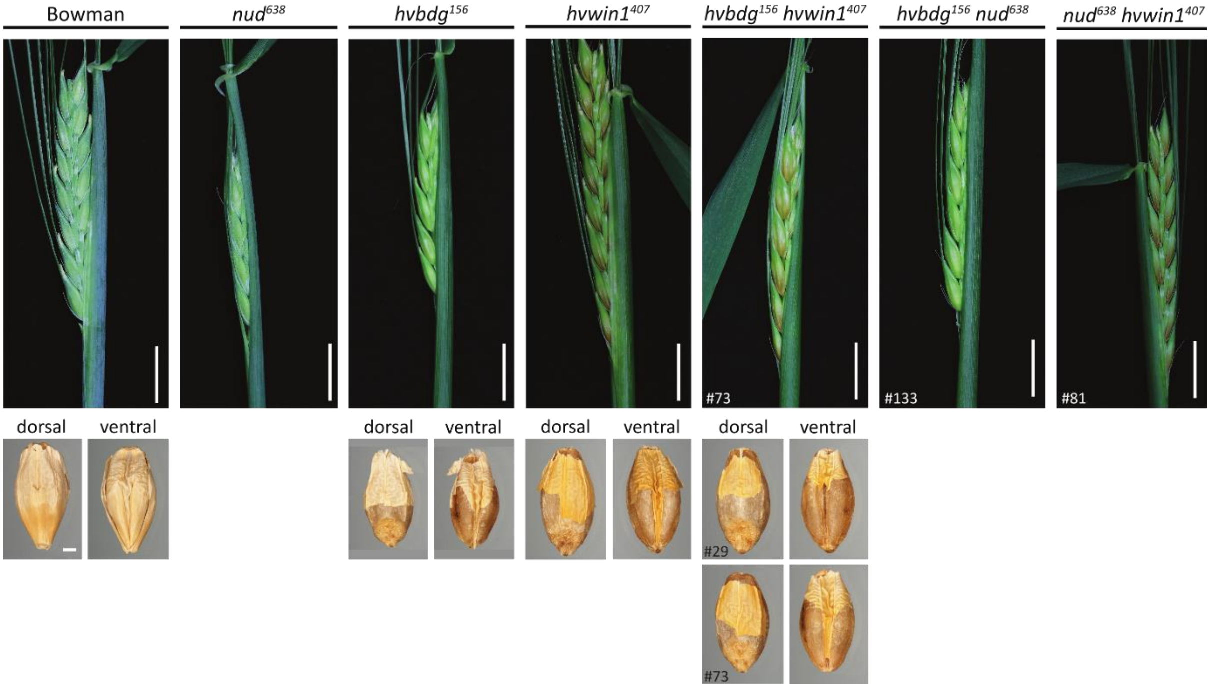


**Fig S10. Single and double mutant phenotypes.** Representative photos of leaf sheaths and hull adhesion in Bowman, single and double mutants. Leaf sheaths shown for Bowman, *nud*$^{638}$*, hvbdg1*$^{156}$, *hvwin1*$^{407}$, *hvbdg1*$^{156}$ *hvwin1*$^{407}$ #73 double mutant, *hvbdg1*$^{156}$ *nud*$^{638}$ #133 double mutant and *nud*$^{638}$ *hvwin1*$^{407}$ #81 double mutant. Scale bar, 1 cm. Manually threshed grain shown for Bowman, *hvbdg1*$^{156}$, *hvwin1*$^{407}$, *hvbdg1*$^{156}$ *hvwin1*$^{407}$ #29 double mutant and *hvbdg1*$^{156}$ *hvwin1*$^{407}$ #73 double mutant. Dorsal and ventral sides. Scale bar, 0.05 cm.

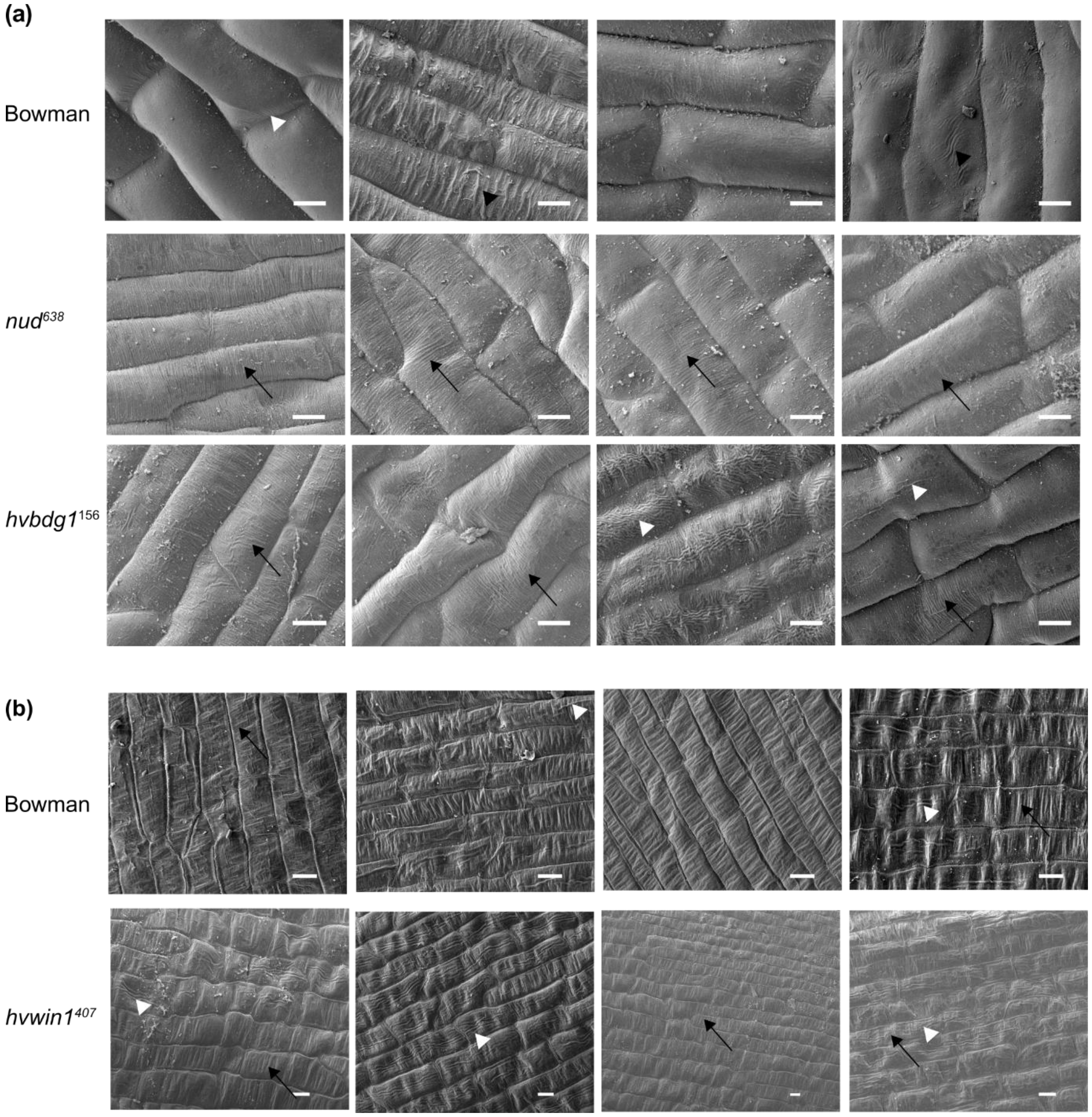


**Fig S11 Cuticular ridges on caryopses**. Scanning electron microscopy (SEM) of 7 days post anthesis (DPA) caryopses (hulls removed) pericarp surfaces. (a) Bowman, *nud*$^{638}$ and *hvbdg1*$^{156}$. (b) Bowman and *hvwin1*$^{407}$ compared in a separate experiment to (a). Scale bars, 10 µm. White arrowheads show longitudinal cuticular ridges, black arrowheads perpendicular wrinkles and black arrows perpendicular nanoridges.

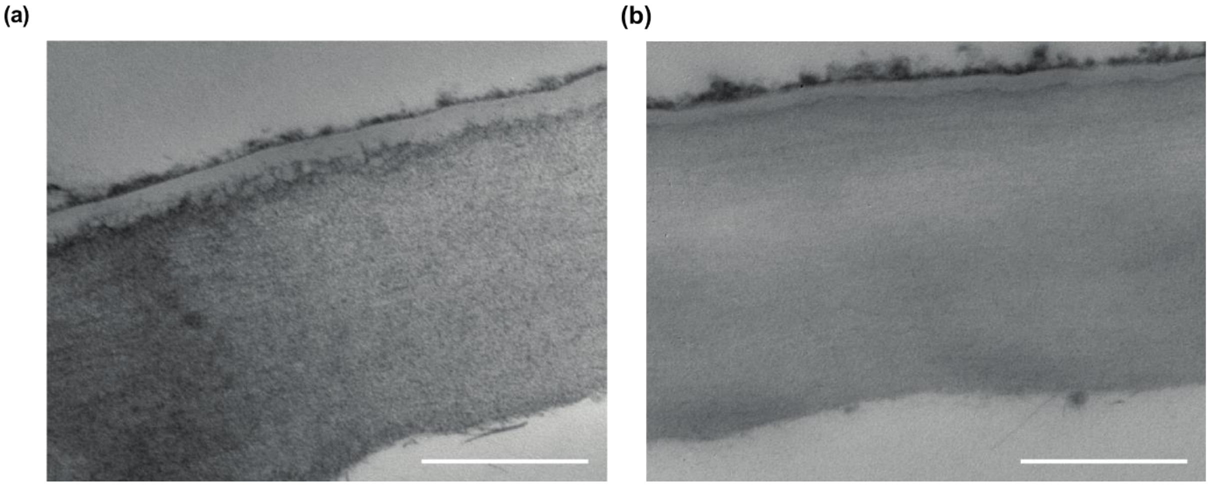


**Fig S12 7 DPA Bowman caryopsis cuticle is thicker at 7 DPA**. (a) Bowman. (b) *nud*[638]. Scale bars, 500 nm. Images are representative of at least three biological replicates.

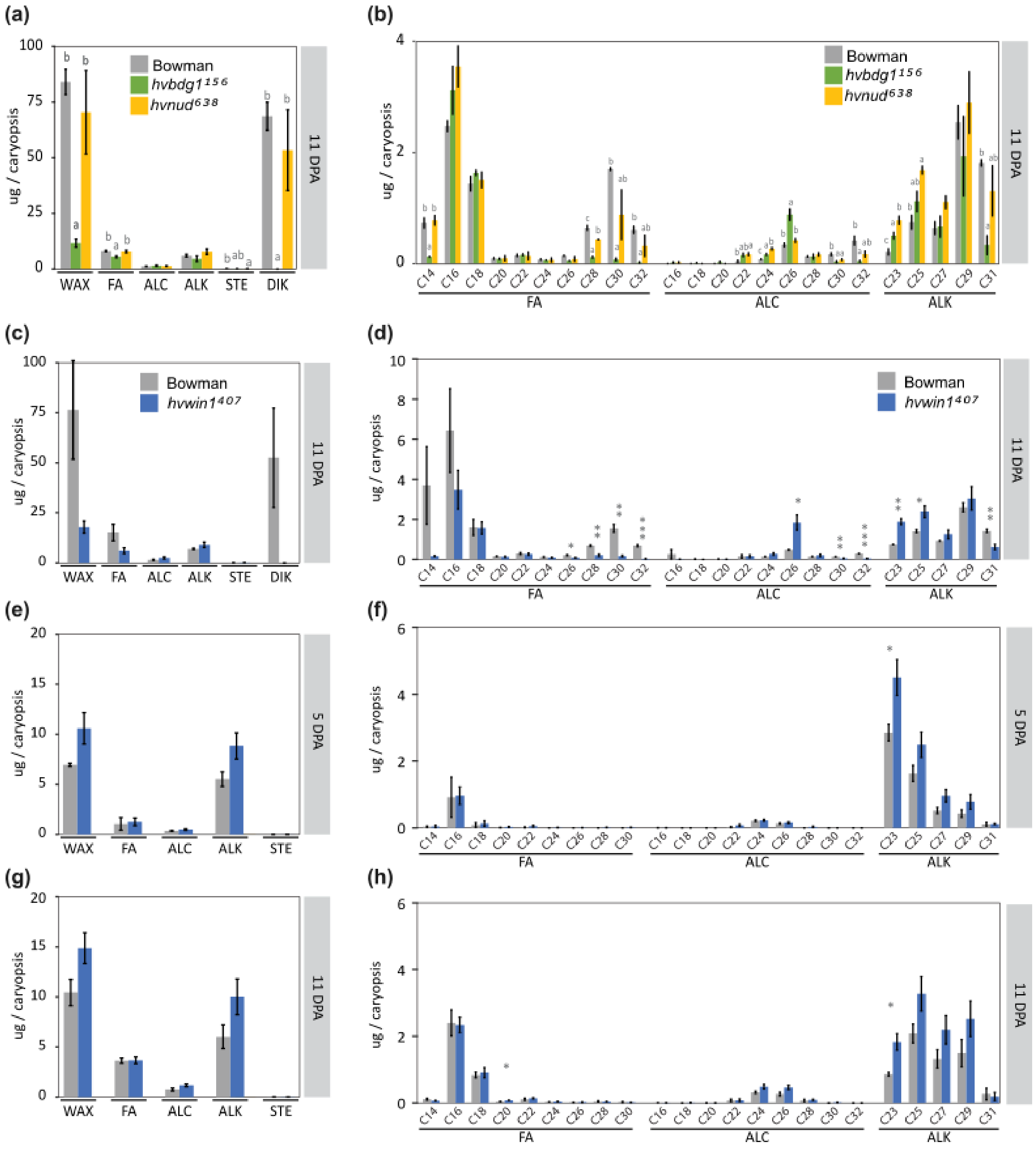


**Fig S13 Surface lipids on hulls and caryopses during adhesion**. (a) Soluble wax classes and (b) Soluble wax chain lengths extracted from hulls dissected from 11 DPA caryopses from Bowman, $nud^{638}$ and $hvbdg1^{156}$. (c) Soluble wax classes and (d) Soluble wax chain lengths extracted from hulls dissected from 11 DPA caryopses from Bowman and $hvwin1^{407}$. (e) Soluble wax classes and (f) Soluble wax chain lengths extracted from 5 DPA caryopses of Bowman and $hvwin1^{407}$. (g) Soluble wax classes and (h) Soluble wax chain lengths extracted from 11 DPA caryopses **of** Bowman and $hvwin1^{407}$. WAX = total wax extracted, FA = fatty acids, ALC = primary alcohols, ALK = alkanes, STE = sterols. FA = fatty acids, ALC = primary alcohols, ALK = alkanes. Bars show the average with standard deviation of three biological replicates. Letters indicate significant differences within genotypes ($P < 0.05$; Tukey's HSD multiple comparison following one-way ANOVA). Asterisks indicate significant differences within genotypes ($P < 0.05$ = *, $P < 0.01$ = **, $P < 0.001$ = ***; one-way ANOVA).

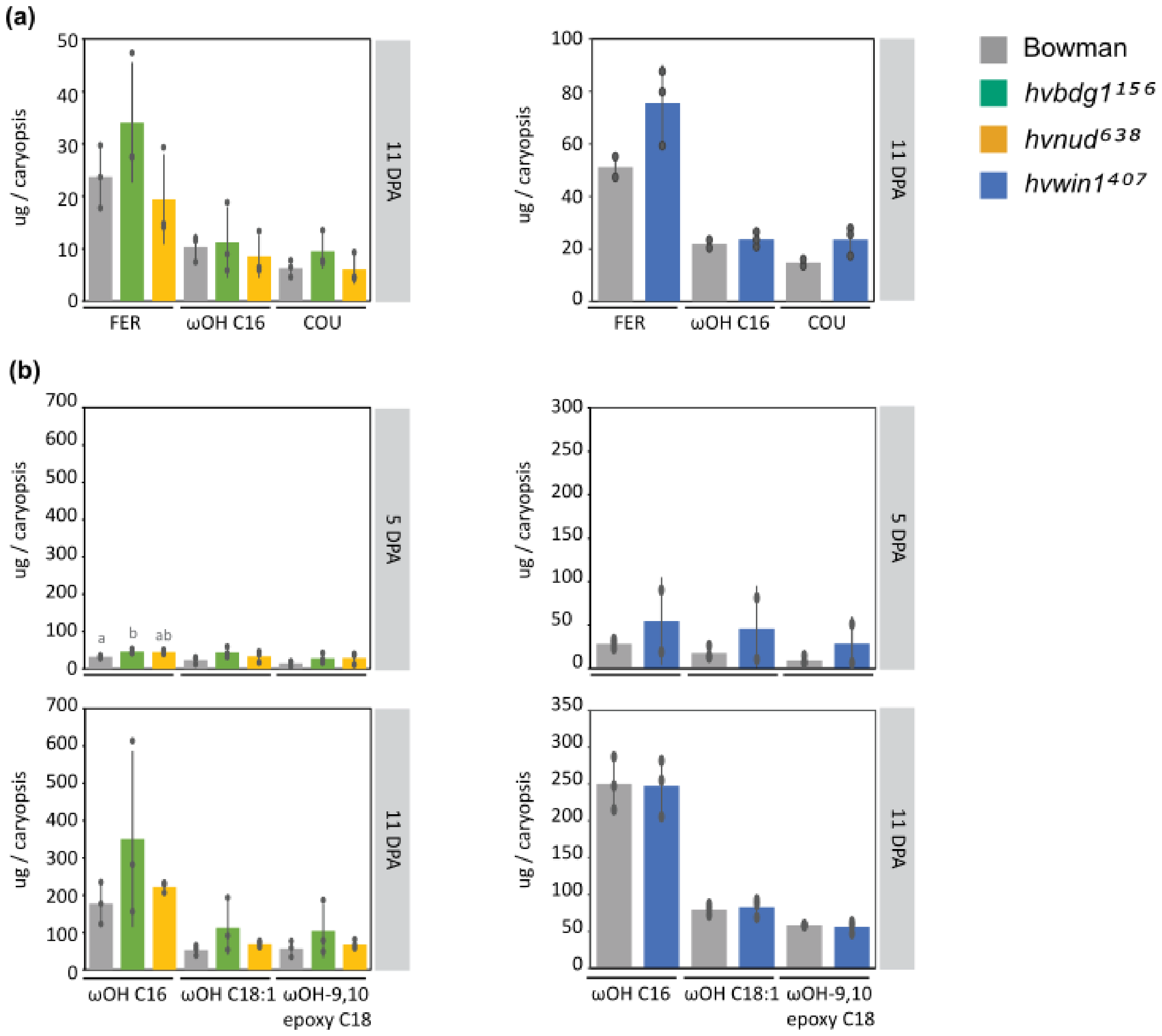


**Fig S14 Cutin monomers from hull and caryopses during adhesion**. (a) Cutin monomers extracted from hulls dissected from 11 DPA hulls from Bowman, *nud*$^{638,}$ *hvbdg1*$^{156}$ and *hvwin1*$^{407}$. (b) Cutin monomers extracted from 5DPA and 11DPA caryopses of Bowman, *nud*$^{638,}$ *hvbdg1*$^{156}$ and *hvwin1*$^{407}$. Bars show the average with standard deviation of three biological replicates, except for Bowman 11DPA hulls in the experiment with *hvwin1*$^{407}$ (right top panel) and *hvwin1*$^{407}$ 5DPA caryopses where replicates were two. Letters indicate significant differences within genotypes ($P < 0.05$; Tukey's HSD multiple comparison following one-way ANOVA).

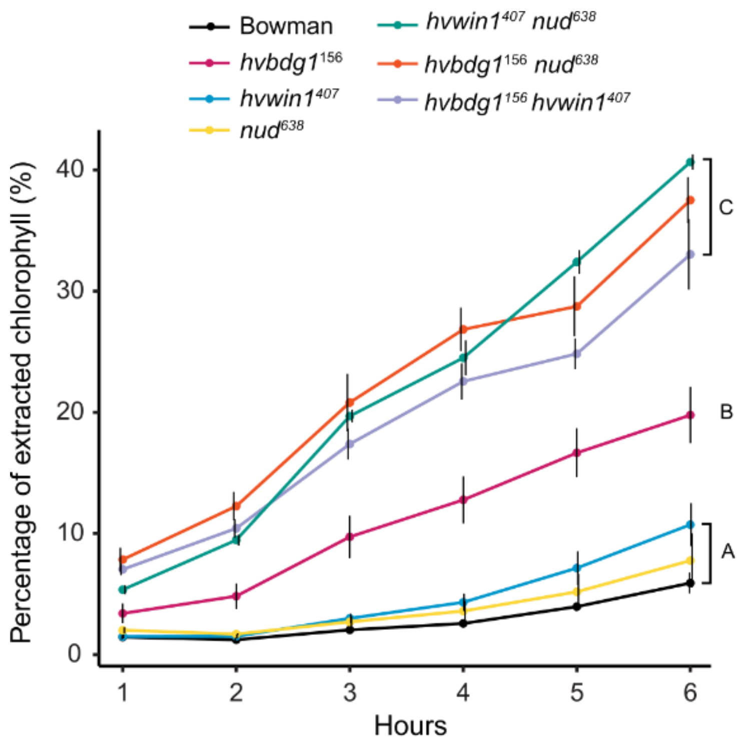


**Fig S15 Chlorophyll leaching in leaf blades**. Chlorophyll leaching of detached leaf blades into 80% (v/v) ethanol. Letters refer to significant differences using an analysis of variance (ANOVA) and the post-hoc Tukey's HSD test using Area Under the Curve (AUC). (n = 4).

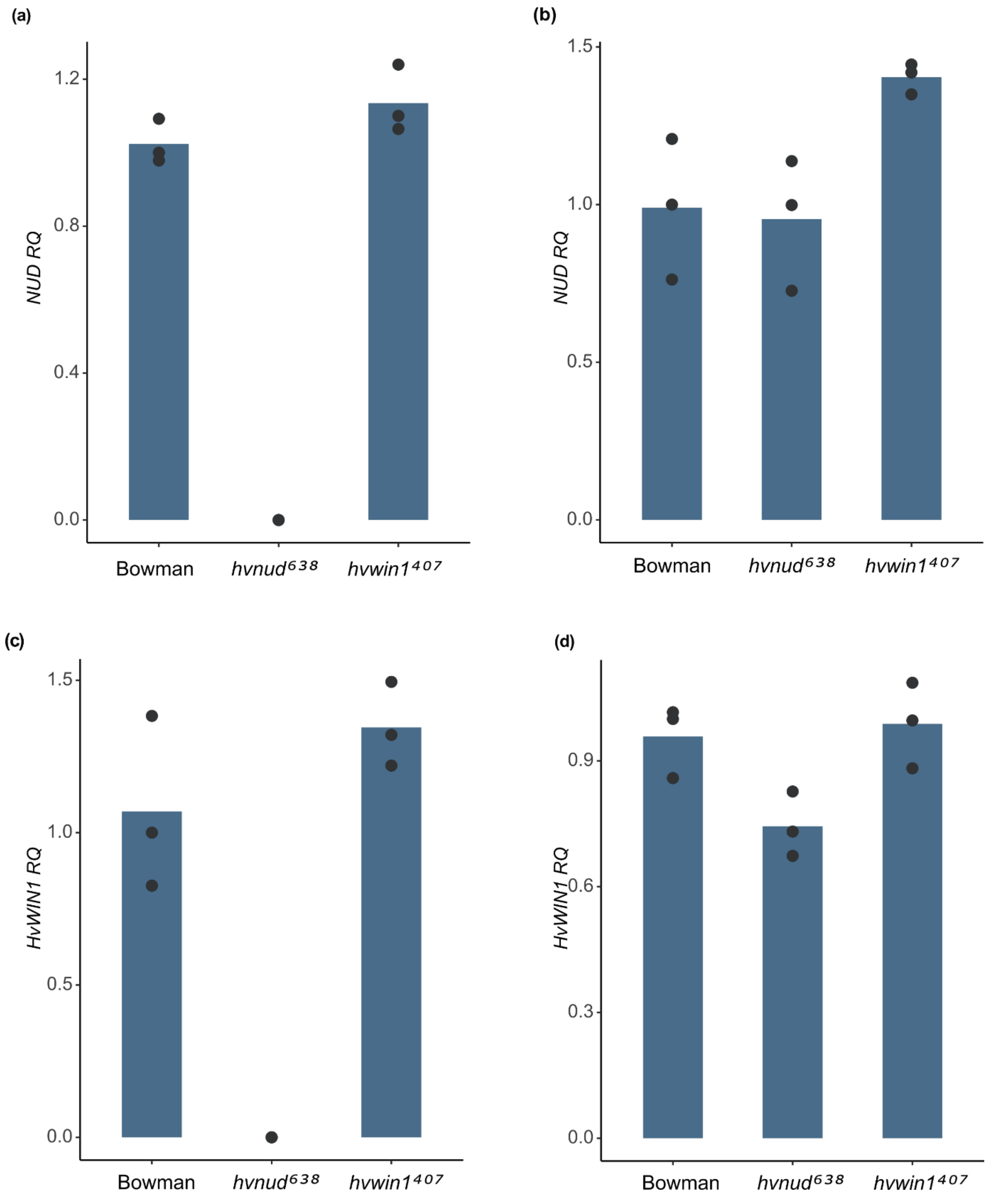


**Fig S16 *NUD* and *HvWIN1* expression in developing second leaf blades**.qRT-PCR of *NUD and HvWIN1* transcript levels in Bowman, *hvnud*$^{638}$ and *hvwin1*$^{407}$ expressed as relative quantity (RQ). (a) *NUD* expression in the leaf base. (b) *HvWIN1* expression in the leaf base. (c) *NUD* expression in the mid-leaf.

(d) *HvWIN1* expression in the mid-leaf. Bars indicate mean expression relative to Bowman. Black circles represent independent biological replicates, each the average of three technical repeats.

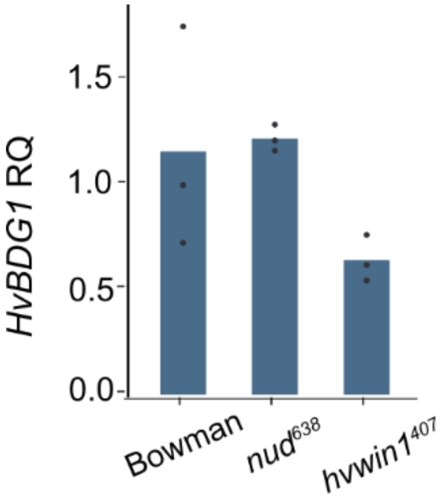


**Fig S17 HvBDG1 expression in midleaf blade sections**. qRT-PCR of *HvBDG1* in Bowman, *hvnud*$^{638}$ and *hvwin1*$^{407}$ midleaf blade sections as expressed as relative quantity (RQ). Bars indicate mean expression relative to Bowman. Black circles represent independent biological replicates, each the average of three technical repeats
